# BayICE: A hierarchical Bayesian deconvolution model with stochastic search variable selection

**DOI:** 10.1101/732743

**Authors:** An-Shun Tai, George C. Tseng, Wen-Ping Hsieh

## Abstract

Gene expression deconvolution is a powerful tool for exploring the microenvironment of complex tissues comprised of multiple cell groups using transcriptomic data. Characterizing cell activities for a particular condition has been regarded as a primary mission against diseases. For example, cancer immunology aims to clarify the role of the immune system in the progression and development of cancer through analyzing the immune cell components of tumors. To that end, many deconvolution methods have been proposed for inferring cell subpopulations within tissues. Nevertheless, two problems limit the practicality of current approaches. First, all approaches use external purified data to preselect cell type-specific genes that contribute to deconvolution. However, some types of cells cannot be found in purified profiles and the genes specifically over- or under-expressed in them cannot be identified. This is particularly a problem in cancer studies. Hence, a preselection strategy that is independent from deconvolution is inappropriate. The second problem is that existing approaches do not recover the expression profiles of unknown cells present in bulk tissues, which results in biased estimation of unknown cell proportions. Furthermore, it causes the shift-invariant property of deconvolution to fail, which then affects the estimation performance. To address these two problems, we propose a novel deconvolution approach, BayICE, which employs hierarchical Bayesian modeling with stochastic search variable selection. We develop a comprehensive Markov chain Monte Carlo procedure through Gibbs sampling to estimate cell proportions, gene expression profiles, and signature genes. Simulation and validation studies illustrate that BayICE outperforms existing deconvolution approaches in estimating cell proportions. Subsequently, we demonstrate an application of BayICE in the RNA sequencing of patients with non-small cell lung cancer. The model is implemented in the R package “BayICE” and the algorithm is available for download.

## 1 Introduction

Exploring the cellular components of heterogeneous tissues from their gene expression profiles is an essential work for revealing molecular mechanisms across different cell types. For instance, increasing evidence suggests that levels of tumor-infiltrating immune cells are associated with tumor progression, response to therapy, and patient survival (Dieu-Nosjean, et al., 2014; Fridman, et al., 2012; Fridman, et al., 2017). Thus, powerful technologies for single-cell isolation, such as laser microdissection and flow cytometry, have been employed to quantify the numbers of malignant and normal cells in tissue (Hu, et al., 2016). However, these physical approaches to isolating cells of interest at the gene expression level are costly and time-consuming, resulting in drastically reduced biological-content yields. In contrast to single-cell technologies, RNA-seq and microarrays yield bulk gene expression from hundreds of thousands of cells. In heterogeneous tissues, where more than one cell type is present, the expression profile from bulk RNA-seq or microarrays is from cell mixtures; thus, to correctly interpret these data, gene expression deconvolution approaches are required to recover cell type-specific expression and the distinct cellular proportions within complex tissues.

In the study of gene expression deconvolution, numerous computational and statistical approaches have been proposed to characterize cell subpopulations within tissues (Anghel, et al., 2015; Becht, et al., 2016; Gong, et al., 2011; Li, et al., 2016; Newman, et al., 2015; Ogundijo and Wang, 2017; Racle, et al., 2017; Xie, et al., 2018; Zhong, et al., 2013). Expression data from a heterogeneous tissue can be modeled as a linear combination of the distinct expression profiles of the cells present in that tissue, weighted by the corresponding cell fractions. These approaches can be grouped into one of three categories depending on whether they use a prior database of cell type-specific expression profiles in the deconvolution procedure: reference-free, reference-based, and semi-reference-based methods. Reference-free approaches aim to directly perform expression deconvolution without cell type-specific references, and their most significant feature is that they estimate the relative cellular proportions and simultaneously disentangle their expression profiles. For instance, many studies have been leveraged on non-negative matrix factorization to decompose mixed gene expression matrices into cell fractions and their corresponding expression profiles (Gaujoux and Seoighe, 2012; Prassas, et al., 2012). Although reference-free models are valuable in the exploration of an uncharacterized cell population, such as tumor subclones (Xie, et al., 2018), relating the cellular components they identify to specific cell types of interest is difficult. Hence, the results of reference-free approaches are unable to clarify the association between a particular cell type and disease progression.

By contrast, reference-based methods incorporate external expression profiles of pure cell samples for deconvolution. For example, the analytical tool CIBERSORT successfully borrows cell type-specific information to predict the immune cell components in blood tissues and tumors through *v*-support vector regression (*v - SVR*) (Charoentong, et al., 2017; Newman, et al., 2015). A fundamental assumption about reference-based models is that all types of cells present in the target tissues are included in the reference set, and the cellular proportions should sum up to one. Unfortunately, the pure expression profile of malignant cells, a key component in tumors, is a great challenge because of the high genetic heterogeneity of tumors. Hence, reference-based models can only derive the relative cell proportions concerning the reference set rather than the exact proportions concerning the microenvironment. Therefore, the relative cell proportions are not comparable across samples. To overcome this problem, TIMER adopts a series of deconvolution procedures to adjust the relative cell proportions with tumor purity, which is the content of malignant cells in a tumor (Li, et al., 2016).

The abovementioned limitation forms the main incentive for developing semi-reference-based deconvolution approaches. In 2017, Racle et al. proposed a framework for estimating the proportion of immune and cancer cells (EPIC) for RNA-seq data (Racle, et al., 2017). EPIC applies least-squares regression with a non-negativity constraint to the deconvolution problem, and requires that the sum of all cell proportions in each tissue must be less than or equal to one. When the sum is not equal to one, one minus the sum of the estimated cell proportions represents the fraction of uncharacterized cells in a tissue that is not accounted for by the reference set; this number is interpreted as the malignant cell proportion in a bulk tumor.

Although semi-reference-based models demonstrate the advantages of incorporating cell-specific information and simultaneously extracting the uncharacterized cell types present in tissues, two problems behind these models should be addressed to complete the framework of gene expression deconvolution. First, signature gene selection is critical to the performance of gene expression deconvolution. In some studies, the incorporation of a preselected signature gene set has successfully improved the accuracy of immune cell deconvolution (Chen, et al., 2017; Newman, et al., 2015; Racle, et al., 2017; Wu, et al., 2018). However, the gene activities of a particular cell type usually vary across different tissue microenvironments. Hence, the use of a preselected gene set might cause the loss of data-dependent information and lead to less deconvolution power. The second problem concerns the natural characteristics of deconvolution. In a reasonable strategy for deconvolution, shifts in the mean level of reference samples and tumor samples should not change the estimation of cell proportions. We refer to this as the shift-invariant property for deconvolution. However, the constrained model implemented for EPIC does not maintain the shift-invariant property in deconvolution, and the estimation of cellular components is unstable because the sequencing depth changes across experiments. More specifically, the uncharacterized cell fractions estimated using the constrained least-squares approach are biased toward zero, which will be demonstrated in our results.

Therefore, to address the aforementioned problems, we propose a new model based on a hierarchical Bayesian framework for intracellular component exploration. It is called BayICE and is a semi-reference-based approach. Under the Gaussian assumption, we first considered stochastic search variable selection (SSVS) for novel signature gene selection (George and McCulloch, 1993). The SSVS approach has been widely used in transcriptome analyses to select significant genes. For example, Ishwaran and Rao introduced a rescaled Bayesian model for selecting differentially expressed genes through multi-group microarray data (Ishwaran and Rao, 2005). To the best of our knowledge, BayICE is the first attempt to incorporate the mechanism of feature selection for inferring the cellular components of bulk tissues. Moreover, we claim that the BayICE model possesses the shift-invariant property of deconvolution, which yields unbiased estimates of cellular proportions. The model with the shift-invariant property further guarantees that it can recover the expression profiles of uncharacterized cells using posterior mean inference. For the purpose of inference, we applied Gibbs sampling and the Metropolis–Hastings as the sampling procedure in the estimation. In brief, BayICE performs cellular component estimation, uncharacterized cell expression profile estimation, and a novel strategy for signature gene selection.

The remainder of this paper is organized as follows. Section 2 introduces the deconvolution in gene expression and states the shift-invariant property. Section 3 introduces the statistical modeling of BayICE for gene expression deconvolution and proposes a Markov chain Monte Carlo (MCMC) algorithm for simulating the posterior distributions of parameters. To assess the model’s performance, Section 4 presents simulation studies that investigate gene expression deconvolution, gene selection, and model robustness compared to two existing methods. Section 5 presents applications to two real datasets where underlying true cell proportions are known and performance can be benchmarked. Section 6 describes application of BayICE to 199 non-small cell lung cancer RNA-seq samples and exploration of the cell components present in the microenvironments of lung tumors. Finally, Section 7 provides final discussion and conclusions.

## 2 Deconvolution

The deconvolution in gene expression can be formalized as an optimization problem in which the parameter of interest is the cellular proportion ***W*** = (*w*_1_ …, *w*_*K*_)′, and the estimates of ***W*** are obtained by

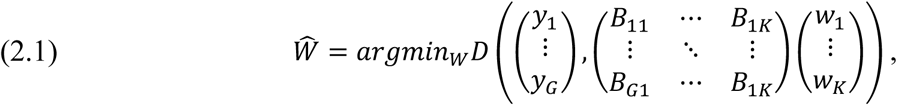

where *y*_*g*_ is the gene expression of gene *g, B*_*gk*_ is the expression of the k-th cell type in gene *g*, and *D* is the distance metric. In general, *D* is the Euclidean distance. As mentioned above, the reference-free deconvolution approaches assume the cell type-specific expression (*B*_*gk*_) is unobserved, and the reference-based methods, by contrast, require *B*_*gk*_ as input for the optimization problem. Additionally, the semi-reference-based approaches allow that one of the cell types is absent in the reference. We now define the shift-invariant property to characterize different deconvolution models.

### Definition 1 (Shift-invariant property)

Let Y be the mixed expression from bulk tissue, and B be the cell type-specific expression matrix. If the cellular proportion estimate of a deconvolution method *M* is invariant when the expression distribution shifts with a constant (location parameter), then *M* has the **shift-invariant property**. That is, *M* satisfies

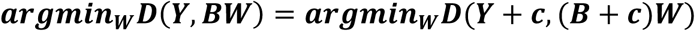

for any constant *c*, where *D* is the distance metric used by *M*.

The shift-invariant property in Definition 1 is essential for evaluating the deconvolution methods, especially in the genomic study. Since the protocol of a gene expression experiment is typically designed for each study, the mean read depth varies across experiments. More specifically, the different experimental protocols cause the location parameters of expression distributions to change. If the location parameter affects the estimation of the same composition, then it is not reasonable to compare results across studies. However, the shift-invariant property guarantees that a deconvolution method with this property can estimate the proportions precisely when the location parameter is changed, and thus, the between-study comparison is valid. In the supplementary file, we have shown that the reference-free and reference-based deconvolution approaches follow the shift-invariant property. By contrast, EPIC adopted the inequality-constrained optimization method to derive cell proportions lacks the shift-invariant property, and hence the following simulation result reveals that the estimates of EPIC are biased.

To address the issue of shift-invariant property for the semi-reference-based deconvolution approaches, we proposed a Bayesian deconvolution model which is more robust to the change in the location parameter. The Bayesian hierarchy architecture facilitates the construction of equality-constrained objective function for the semi-reference-based deconvolution problem via likelihood approach. The proof details in the supplementary material, and the model construction will be detailed in the next section.

## 3 BayICE Deconvolution Model

In this section, we present the proposed hierarchical Bayesian deconvolution model for intracellular component exploration with novel signature gene selection. Figure 1 provides a graphical representation of the BayICE hierarchical model. We first describe the input data and establish the statistical modeling for reference samples and tumor samples. Subsequently, a stochastic search method, the Bayesian false discovery rate, and an inflation factor are introduced for signature gene selection. Finally, we adopt the Gibbs sampling approach and the Metropolis–Hastings approach to develop a comprehensive sampling procedure.

**Figure 1.**
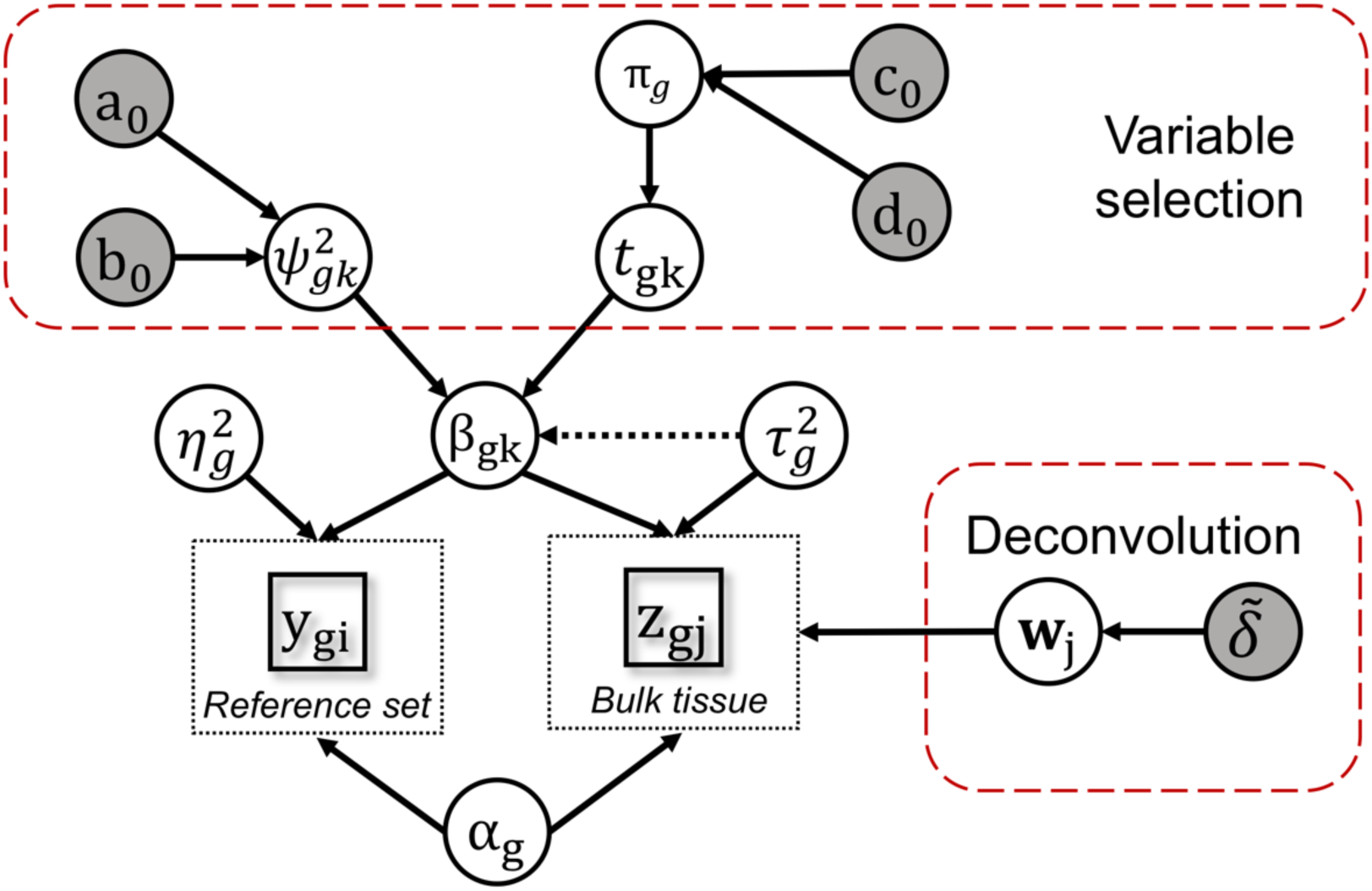
Bayesian hierarchical model of BayICE. The square symbols represent observed data from the reference set and bulk tissues. The white circles indicate priors and the grey circles are hyperparameters.

### 3.1 Input data and normalization

BayICE is a statistical framework designed for gene expression deconvolution. We first assume that the gene expression data are available from two sets of samples, namely heterogeneous tissues with a given clinical condition and a reference set consisting of several groups of samples with pure cell types. Although BayICE is a quantitative model-based approach, microarray and RNA-seq expression data are both legal inputs for BayICE. For the microarray data, we implement a locally weighted scatterplot smoothing algorithm (Yang, et al., 2002) for normalization. For the read count RNA-seq data, we recommend two different strategies for normalization. The first is to use new gene expression units, called transcripts per million (TPMs), which were proposed by Wagner et al. in 2012. The main feature of TPM normalization is to make the sum of all TPMs equal across samples, which facilitates fair comparisons between samples. For public data, TPM normalization approach is not always available because the information of sequencing depth or gene length required for TPM could be missing. Thus, we adopt subsampling normalization as the second strategy. Subsampling normalization applies a binomial sampler to resample the read count, and it aims to maintain internal associations between genes and can simultaneously adjust external variance between samples. The details of how we implement the two normalization approaches are provided in the supplementary materials. After normalization by the first strategy or the second strategy, we consider a log transformation by log(count + 1). The log transformation of count data has been widely applied in RNA-seq studies.

### 3.2 Statistical modeling

The problem of gene expression deconvolution can be formulated as a system of linear equations that describes the expression of a given gene in a bulk tissue as the weighted sum of the expression values from multiple cell types present in the tissue. To maximize the deconvolution power, BayICE incorporates a reference set comprising cell-specific expression profiles into the inference of cellular components in bulk tissues. In this study, the reference set contains various types of immune cells, such as T cells, B cells, natural killer cells, monocytes, neutrophils, and normal tissue cells. In addition to these nonmalignant cells, malignant cells are a major cellular component of tissues in cancer deconvolution studies. Unfortunately, the high genetic heterogeneity of malignant cells hinders the possibility of constructing a predefined cancerous cell expression profile that can be applied to every bulk sample. To address this problem, BayICE takes the advantages of the flexibility of hierarchical Bayesian inference to extract malignant cell profiles directly from bulk tissues. Moreover, BayICE infers cellular proportions with respect to the reference cell types and unknown cell types of each tissue by integrating with a Bayesian signature gene selection approach.

We first introduce the statistical model for the reference set consisting of N purified samples with K cell types. We denote an observation in this set as *y*_*gi*_, which is the normalized value for gene-*g* (*g* = 1, …, G) in the *i*-th purified sample (*i* = 1, …, N). The N purified samples belong to K nonmalignant cell types that exist in the bulk tissues. We introduce a binary vector variable *x*_*i*._ = (*x*_*i*1_, …, *x*_*iK*_) to represent the cell type for the *i*-th purified sample. More specifically, if the *i*-th purified sample belongs to cell type *k*, then *x*_*ik*_ = 1 and *x*_*it*_ = 0 for all *t* ≠ *k*. BayICE assumes that *y*_*gi*_ follows the Gaussian distribution with a mean level *μ*_*gi*_ and a gene-specific variance denoted by 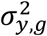. For the purpose of signature gene identification, we consider the mean structure *μ*_*gi*_ to comprise a gene-specific baseline (*α*_*g*_) and cell type-specific effects (*β*_*gk*_). Thus, the modeling of reference data can be written as

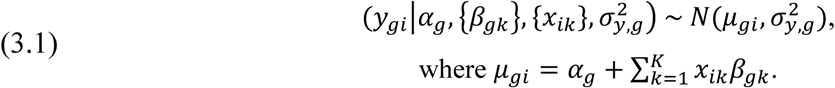

To search for signature genes that exhibit a differential effect among cell types, we define a baseline cell type against which changes in expression levels are measured. We set *β*_*g*1_ to be 0 for all gene-*g*.

The cell type-specific effects (*β*_*g*1_, …, *β*_*gK*_) in (3.1) are shared in constructing the mean structure of gene expression in bulk tissues. It has been observed in the literature that gene expression of a certain cell type changed when it went through cell sorting (Richardson, et al., 2015; van den Brink, et al., 2017). To accommodate the effect of the changes induced by cell separation, we adopt the joint modeling of *β*_*g*1_, …, *β*_*gK*_ for pure cell samples and bulk samples.

For the model of the bulk tissue, it is a linear combination of K nonmalignant cell-specific expression profiles and malignant cell gene expression. The expression of gene-*g* from the *j*-th bulk tissue (*j* = 1, …, M) is denoted by *z*_*gj*_. The mean of *z*_*gj*_, called *v*_*gj*_, can be modeled as the weighted sum of nonmalignant cell-specific effects {*β*_*gk*_} and one malignant cell-specific effect *β*_*g*0_ through a linear regression, given by

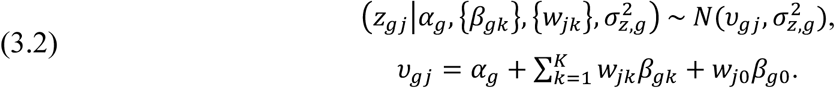

The weights, *w*_*jk*_, are the proportions of expression attributable to normal cell type k in the *j*-th tumor, and the weight, *w*_*j*0_, represents tumor purity, which is the percentage of malignant cells in a tumor tissue. Notably, BayICE can be used to explore not only tumors but also other noncancerous tissues. For noncancerous tissues, *w*_*j*0_ can refer to the proportion of one unknown cell type that is uncharacterized by the reference set. In our model, a natural constraint for these cell proportions is that the sum of weights across cell types should be one (i.e., 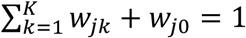 for each *j*). Furthermore, to characterize the gene expression pattern in malignant cells, we introduce a tumor-specific parameter *β*_*g*0_ to represent the effect size of gene-*g* in cancer. In a real application, multiple unknown cell types or cancerous cell types might be present in a bulk tissue. In this case, BayICE treats these uncharacterized cell types as a whole, and *w*_*j*0_ represents the proportion of unknown class in a bulk tissue.

### 3.3 Novel gene selection using the SSVS approach

Identifying signature genes that are expressed in a particular cell type is essential to the success of expression deconvolution. Although a preselected signature gene set could be easily applied to data analysis, the application of external signature genes could lose data-dependent information for deconvolution. Thus, we incorporate the stochastic search variable selection (SSVS) approach into our Bayesian deconvolution model for integrating expression deconvolution with novel signature gene selection. The SSVS approach, introduced by George and McCulloch (1993), specifies a spike-and-slab mixture prior, which uses data to extract the potential features of the true model by inferring posterior probability. The spike component, which concentrates its mass at values close to zero, shrinks small effects to zero, whereas the slab component spreads its mass over a wide range of possible values for the effect size.

The proposed prior structure of BayICE on effect size exhibits a bimodal distribution on the variance of the coefficients that result in a spike-and-slab type prior on the effects themselves (Ishwaran, et al., 2010; Ishwaran and Rao, 2005). For each effect size *β*_*gk*_, the prior structure is given by

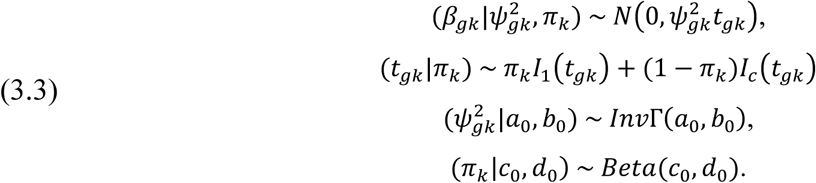

where *I*_c_(*t*_*gk*_) is a mass function that is 1 at *t*_*gk*_ =*c* and 0 everywhere else. We set the value c as a small positive number in this study, such as *c* = 10^−5^, and thus the random variable *t*_*gk*_ is 1 with probability *π*_*k*_ and close to zero with probability 1 − *π*_*k*_. When the *g*-th gene is differentially expressed between the k-th cell type and the other types, *β*_*gk*_ is more likely generated from the slab component and *t*_*gk*_ equals one. By contrast, *t*_*gk*_ = *c* indicates that the *g*-th gene is irrelevant to cell types and its effect size is from the spike component. The hypervariance 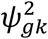 is sampled from an inverse gamma with two given hyperparameters, *a*_0_ and *b*_0_. Following Ishwaran and Rao (2005), *a*_0_ and *b*_0_ are set as 5 and 50, respectively. The proportion of genes differentially expressed in cell type k is controlled by *π*_*k*_, and we assume that *π*_*k*_ follows a beta distribution with *c*_0_ = 0.1 and *d*_0_ = 0.1.

To borrow variance information across samples, we modify the variance structure of effect size *β*_*gk*_ in (3.3) by considering the gene-specific variance 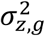 of mixture samples in (3.2) and the modified prior structure for *β*_*gk*_ is given by

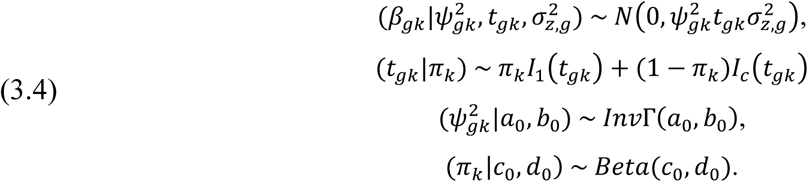

The role of the gene-specific variance 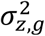 that appears in (3.4) can be intuitively interpreted as the baseline in the feature selection procedure. The modified prior structure considers the trade-off between the value of effect size and gene-specific variance to facilitate the establishment of feature selection.

### 3.4 Bayesian false discovery rate

In frequentist approaches to the test multiplicity problem, controlling the false discovery rate (FDR) has been widely applied to more adequately control genome-wide false positives. Whittemore in 2007 introduced a Bayesian FDR associated analogously with the frequentist FDR (Whittemore, 2007) as follows:

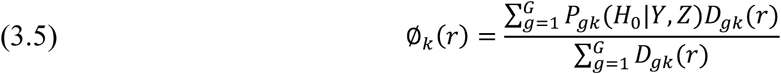

where *P*_*gk*_(*H*_0_|*Y, Z*) is the posterior probability that gene-*g* is not associated with cell type k (H_0_) given observation (Y, Z), and *D*_*gk*_(*r*) is the rejection rule defined by I(*P*_*gk*_(*H*_0_|*Y, Z*) < *r*). The tuning parameter r can be adjusted to control the Bayesian FDR at a certain α level. In the following simulations and applications, the Bayesian FDR is used to address the multiplicity problem.

### 3.5 Inflation factor

In a Bayesian framework, the influence of priors on posterior always vanishes as the sample size increases. This phenomenon limits Bayesian variable selection because the mechanism of such selection requires an effective prior setting. To overcome this disadvantage, Ishwaran and Rao (2005) proposed a rescaling approach to enable estimation invariant to the sample size by setting the prior as a function of the sample size. They applied data rescaling to the gene selection framework for multi-group microarray data. Furthermore, to achieve invariance to sample size, Ishwaran and Rao performed a sample size-related transformation of gene expression through multiplication by the global inflation factor

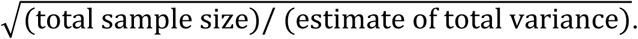

Multiplication by this global inflation factor has been shown to ensure that the prior has a nonvanishing effect. Hence, following the concept of data transformation, we rescale the observations in our reference set and bulk tissue set as follows:

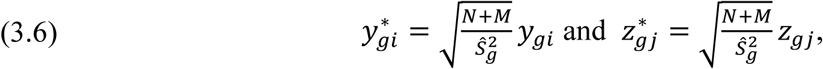

where 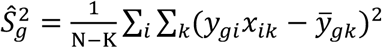 is an unbiased estimator of 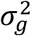 calculated from the reference set. Although we assume that the variance 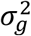 of reference data {*y*_*gi*_} is shared with tissue data {*z*_*gj*_}, the calculation of an unbiased estimator using both {*y*_*gi*_} and {*z*_*gj*_} data is a difficult task because of the convolution structure in {*z*_*gj*_}. Note that the multiplier in (3.6) is a gene-specific inflation factor rather than the abovementioned global inflation factor since the inflation factor is composed of the total sample size and a gene-related variance. Therefore, the use of gene-specific factors can simultaneously achieve sample size invariance and gene-scale consistency.

After rescaling, the corresponding distributions for the transformed data 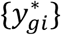 and 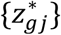 are modified as follows:

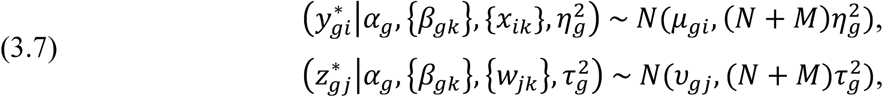

The new variances of the transformed data are adjusted as sample size-related parameters, and this adjustment can be interpreted as a penalization shrinkage effect of the posterior mean. After the adjustment with inflation factors, the variances of 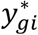 and 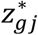 are asymptotically equal to N+M. For the purpose of variable selection, we introduce two further parameters 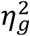 and 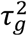 for variances in (3.7) to keep the flexibility of modeling.

### 3.6 MCMC sampling procedure

Next, based on the transformed data 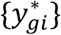 and 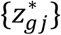, we complete the structure of the hierarchical Bayesian model in BayICE and then establish a sampling procedure to achieve signature gene selection and cell component inference. Following the abovementioned specification, the BayICE model is given by

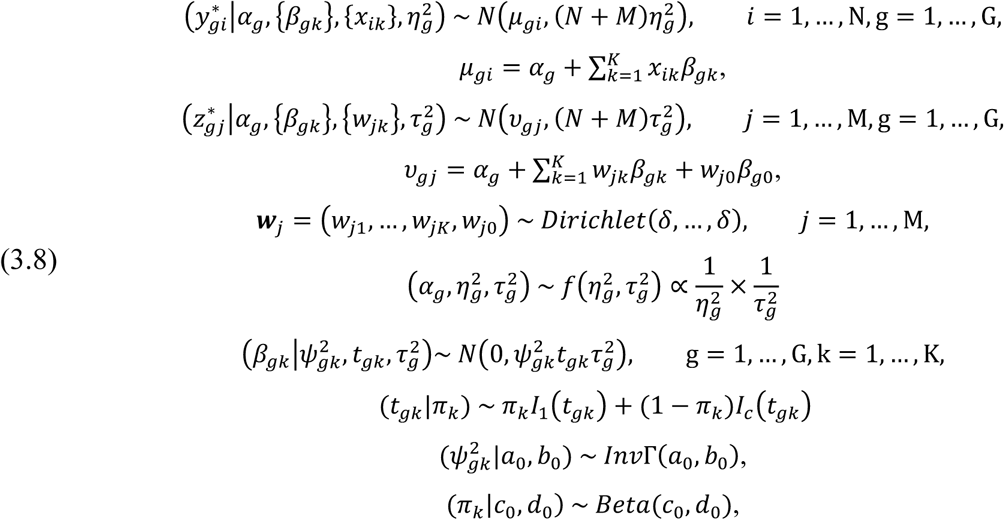

where hyperparameters are specified as *a*_0_ = 5, *b*_0_ = 50, *c*_0_ = 1, *d*_0_ = 1, and δ = 1 in this study.

Subsequently, based on the hierarchical prior setting, we apply Gibbs sampling and the Metropolis–Hastings approach to simulate the posterior value from

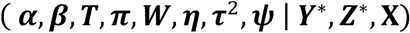

where 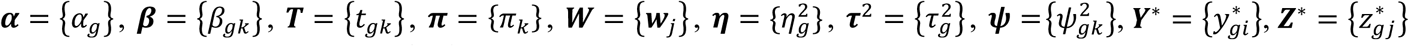, and **X** ={*x*_*ik*_}. The Gibbs sampler in BayICE works are shown in the supplementary file. After a number of iterations, we specify the burn-in period and the thinning interval to obtain the posterior distribution of the parameters and then perform an analysis using the posterior mean or posterior median. In BayICE, the default length of burn-in is 0.6 times the number of iterations, and the thinning interval has a length of 3 iterations. The convergence property is discussed in the supplementary materials.

## 4 Simulation Study

This section presents simulation results based on synthetic datasets to benchmark the performance of BayICE. We consider two types of data: array-based data, which is generated by a normal simulator, and sequencing-based data, which can be produced by a multinomial simulator or negative binomial simulator. We mainly use the multinomial simulator to demonstrate the estimation of cellular proportions, detection of signature genes, and the ability to recover the expression profile of unknown or malignant cells. Furthermore, we apply three different simulators to demonstrate the robustness of BayICE, and the results of the robustness study are shown in the supplementary file.

### 4.1 Multinomial simulator settings

We first perform a sequencing-based simulation using a multinomial simulator for expression deconvolution. Thus, we consider a scenario in which five distinct cell subpopulations are present in a tissue, and one cell type is absent from our reference set. This simulation includes 5000 genes, with 300 genes designed as cell type-related genes. These 300 genes are divided randomly into five disjoint groups, and the genes assigned to a particular group are associated with one cell type. For each cell type in a reference set, we have 20 replicates; hence, the reference set includes 80 samples. In the reference set, an 80 × 4 matrix of binary variables X = {*x*_*ik*_} is used to record the cell type of samples. Furthermore, in the bulk dataset, we simulate the expression data of 90 mixed samples with different cell proportions. The simulation procedure is described as follows.

To account for the fact that the expression level varies across genes, we simulated data according to a set of real data from purified samples. We collected 19 RNA-seq samples of normal lung tissues from the Gene Expression Omnibus database with accession number GSE81089 (Mezheyeuski, et al., 2018), and then took the average gene expression across 19 samples to obtain the baseline 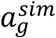 of the g-th gene. Among 17,775 genes, we randomly picked 5000 genes for our simulation study. Based on each gene-specific expression level, we define the cell-specific effect as follows:

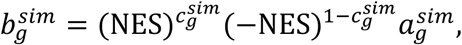

where 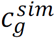 is a binary variable, which is 1 for upregulated status and 0 for downregulated status, and NES is the normalized effect size set as one of the numbers {0.1, 0.2, …, 0.6} The number 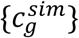 is randomly generated with a probability of 0.5 for 0 and 1. We further sample a series of values from Uniform(0.9,1.1), called 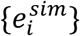, as sample-specific effects because of the sample heterogeneity. Finally, we generate cellular proportions 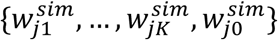. To explore the effect of unknown cell content on the model performance, we assign a fixed number to 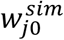, and the remaining components 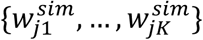 are generated from the Dirichlet distribution with parameters (1,…,1). Because of the sum-to-one constraint on cellular proportions, 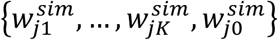 should be normalized by dividing their sum.

The mean expression structure is determined for both purified and mixed samples according to the abovementioned parameter settings. Notably, the use of a multinomial model in our simulation is to simulate the sequence alignment procedure that maps the reads against the reference sequence. The number of trials in the multinomial model is related to the total reads, and we set it as 5 × 10^6^; in other words, the average read depth is designed as 1000. The probability of the multinomial model controls the expression levels across genes, and therefore, we use the relative mean expression as the probability value. The complete sampling procedure details in the supplementary file.

### 4.2 Assessing the inference of deconvolution

To assess the performance of BayICE deconvolution, we include two semi-reference-based approaches for comparison: EPIC and non-negative least-squares (NNLS). NNLS is a general approach for solving the constrained least-squares problem where the coefficients are not allowed to become negative. We modify the NNLS approach by restricting the sum of coefficients to less than one for incomplete reference data deconvolution. We first examine the cellular proportion estimations obtained using EPIC, NNLS, and BayICE. Notably, EPIC and NNLS both require an external step to identify signature genes before deconvolution. In this case, we applied the marker genes identified by BayICE into EPIC and NNLS for a fair comparison.

According to the simulation setting, we generate 90 bulk samples per simulation in which the unknown cell proportions vary from 0.1 to 0.9. A large unknown cell proportion value indicates that the corresponding tissue is highly heterogeneous, such as in tumors; by contrast, a small proportion simulates the microenvironment of normal tissues in which most of the cell types can be purified. Additionally, we evaluate the performance of each model under different normalized effect sizes of marker genes. Moreover, BayICE can recover the underlying expression profile of the unknown cell type using the posterior mean. Because EPIC and NNLS cannot infer unknown expression profiles, we directly compare the estimation of uncharacterized cell profiles with the true mean expression. The results are shown in Figures 2 and 3.

**Figure 2.**
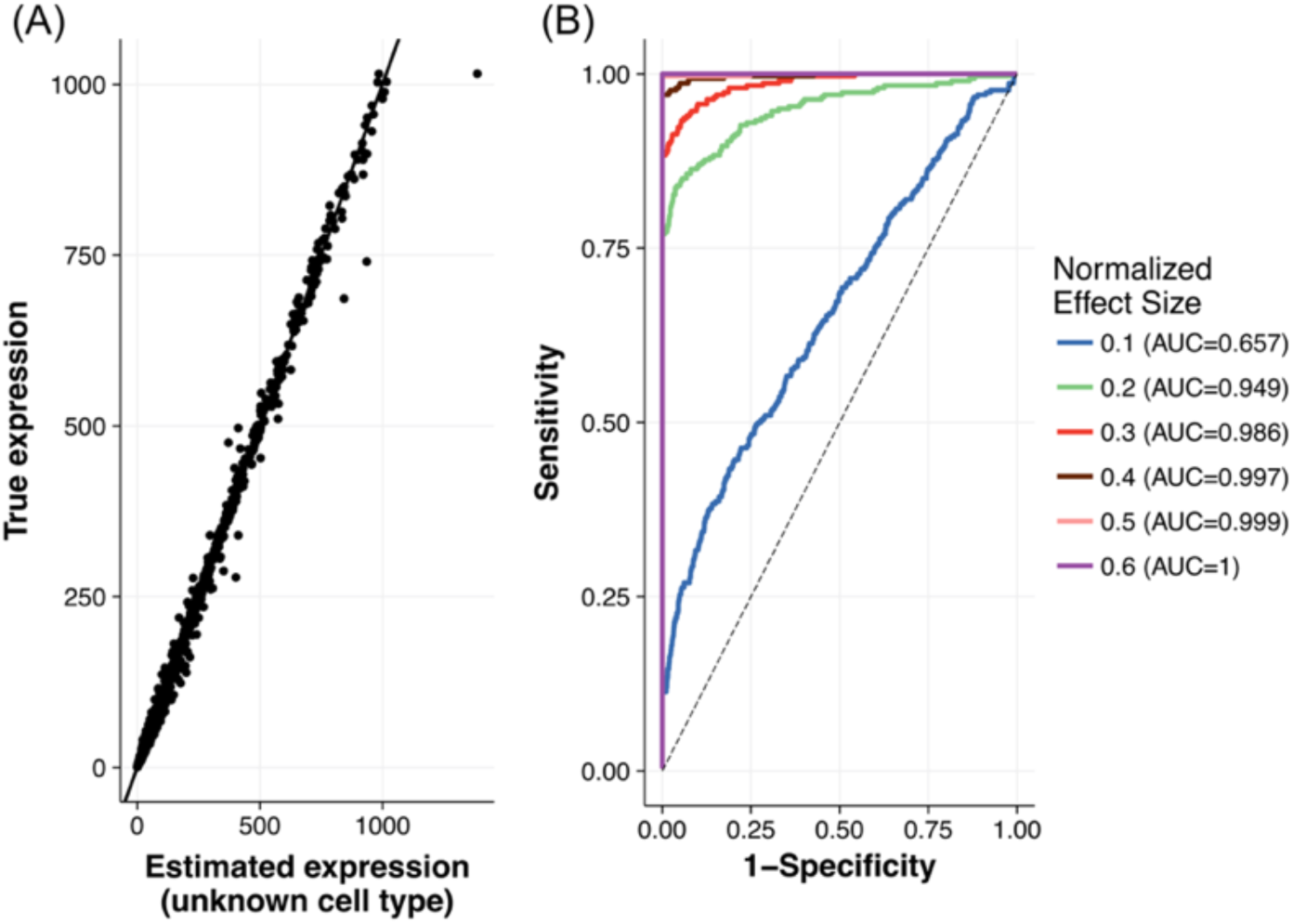
Results of unknown profile estimation and marker gene selection. (A) Scatter plot of gene expression of the unknown cell type between the truth and estimation. (B) ROC curves to evaluate the gene selection of BayICE under different normalized effect sizes.

**Figure 3.**
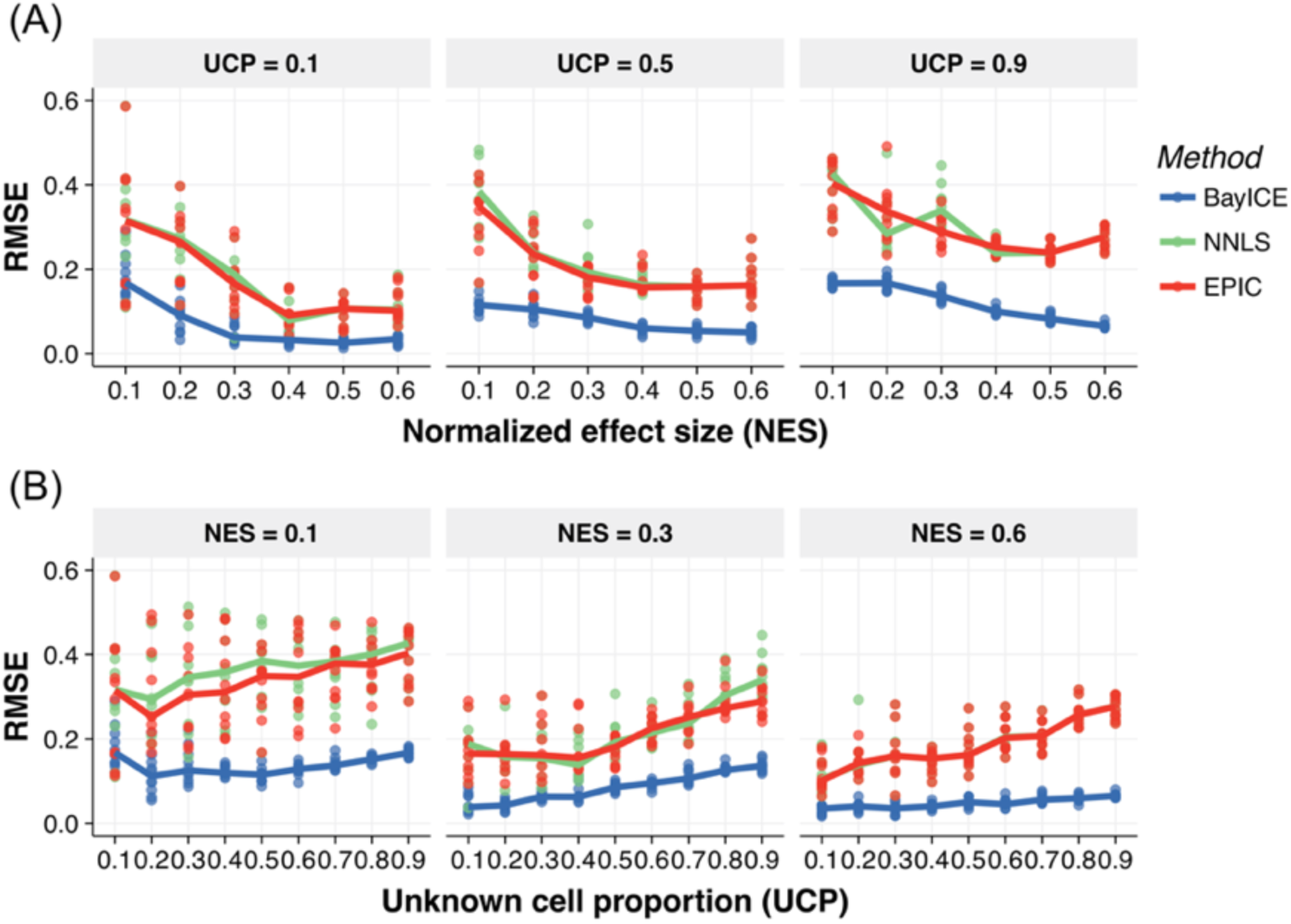
Deconvolution results of cell proportion estimation. Root-mean-square error (RMSE) between the true and estimated cellular proportions under (A) different effect sizes or (B) different proportions of unknown cells. Ten random sets were generated for each condition, and the RMSE in a set was calculated with five pairs of numbers. The medians of the 10 random sets were connected as the line in the figure.

We first evaluate the estimation of gene expression profiles for the unknown cell type and gene identification from BayICE in Figure 2. Figure 2(A) is a scatter plot between the true expression of the unknown cell type and the estimated expression. The results reveal the correlation between the real and estimated values to be greater than 0.98, which implies that BayICE can recover uncharacterized cell expression when one cell type is absent from the reference set. In Figure 2(B), we adopt the receiver operating characteristic (ROC) curve to quantify the results of gene identification under different normalized effect sizes which represent the strength of cell type-specific activity in marker genes. The performance in terms of area under the curve (AUC) is significantly improved with the increase of the effect size, and more specifically, the AUC exceeds 0.94 when the effect is larger than 0.2. Subsequently, we apply root-mean-square error (RMSE) to quantify the accuracy of cell proportion estimation from BayICE, EPIC, and NNLS. Figure 3(A) illustrates the changes in RMSE of all estimates with the increase of normalized effect size under the settings of unknown cell proportion = 0.1, 0.5, and 0.9. In Figure 3(B), we fix the normalized effect size at 0.1, 0.3, and 0.6 and then evaluate those approaches across different proportions of unknown cells. As a result, it is clear that the performance of all approaches decreases when the unknown cell content increases or the effect size decreases. The abovementioned phenomenon reflects that these gene expression deconvolution approaches are less stable in their inference for tissues with highly uncharacterized content or weak cell type-specific signal. However, the simulation shows that the effects of high unknown cell content and low effect size on BayICE estimation are less severe, and overall, BayICE outperforms the other methods.

### 4.3 Evaluating gene detection accuracy

The spike-and-slab prior in our model provides a natural consequence of gene selection. In contrast, the existing models require an external tool to select differentially expressed genes before deconvolution, and DESeq and edgeR are two popular gene selection tools adopted by the deconvolution models. To evaluate the accuracy of gene detection, we compare with DESeq and edgeR. It is worth noted that the genes specifically expressed in the unknown cell type cannot be identified using DESeq and edgeR, and hence the unknown cell type-related genes are excluded in the evaluation. According to the simulation setting, each gene set is differentially expressed in one cell type. For simplicity, the effect size of marker genes is fixed at 0.2, and the cell proportions are randomly decided across samples. In this simulation study, we also assume some genes express inconsistently between purified cells and the same cell type in bulk samples. For each cell type, we randomly select 100 genes from the gene pool to be inconsistent genes, and disturbed the expression of these inconsistent genes by multiplying an inconsistency level. The values of inconsistency level are 0.1, 0.2, 0.3, 0.4, and 0.5, and the mean of the disturbed gene expression is the original mean in Section 4.1 multiplied by 0.9, 0.8, 0.7, 0.6, or 0.5.

In this partial comparison, we evaluate the accuracy of identifying genes related to the four cell types present in the reference. AUC is used to assess the gene detection under different levels of inconsistency, and the results are shown in Figure 4. For the low inconsistency level, the AUCs among three methods are close, and it implies that BayICE is comparable to the other approaches designed explicitly for gene selection. In the case of the considerable inconsistency in gene expression, the AUC of BayICE outperforms DESeqs2 and edgeR, and it shows that BayICE succeeds in borrowing information from mixed bulk samples for gene detection. Consequently, this partial comparison reveals that BayICE can efficiently recover the findings of the two-step approaches when cell activity causes the difference in gene expression between pure cells and tumors.

**Figure 4.**
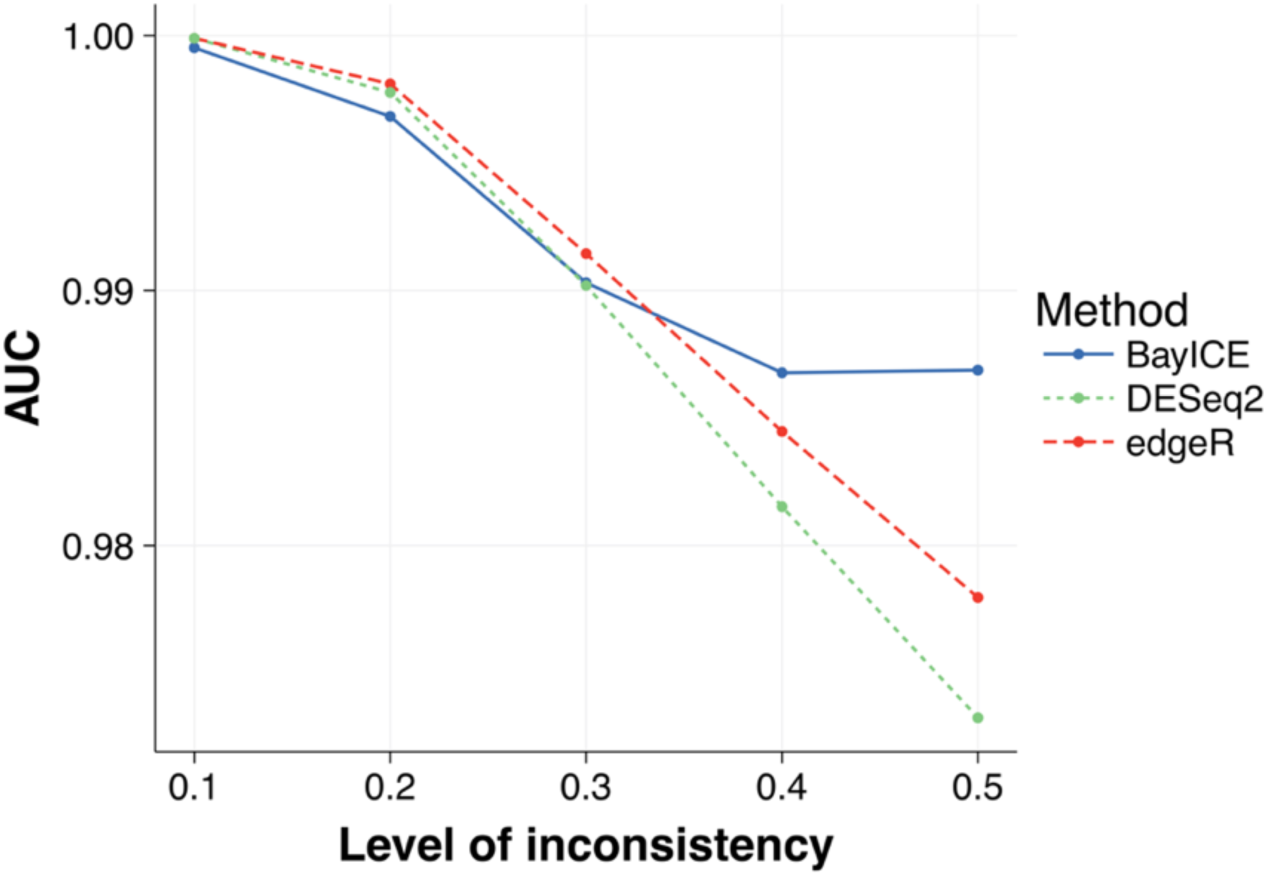
AUC for gene identification. Comparison of the AUC performance on gene identification. Measuring the performances of BayIC, DESeq2, and edgeR at identifying marker genes as measured by the Area Under the ROC Curve (AUC).

## 5 Validation in real data with true proportions

In this section, we consider two microarray mixture experiments for validation. They can be downloaded from the Gene Expression Omnibus database using the accession numbers GSE19830 and GSE11058 (Abbas, et al., 2009; Shen-Orr, et al., 2010). The samples from GSE19830 were mixed with rat brain, liver, and lung tissue derived from the same animal in different proportions. The samples in GSE11058 were mixed with four immune cell lines, Jurkat, IM-9, Raji, and THP-1 at various proportions. To validate the performance in the case of the incomplete reference set, we masked one cell type from the reference set and treated the excluded type as the unknown cell component in tissues. Although EPIC is designed for RNA-seq data, the core concept of EPIC modeling is constrained least-squares optimization, and it can be widely applied to various types of data. Hence, we also performed comparisons with EPIC and NNLS on the abovementioned two real datasets.

Several microarray studies have confirmed that deconvolution on raw-scale data rather than on log-scale data can accurately reflect the underlying cellular components. The raw scale of the data was adopted in this validation study. The signature gene selection for the methods we compared in the microarray followed Hunt et al. (Hunt, et al., 2018), using a *t*-test between one cell type and all other cell types for each gene, and we selected the top 200 differentially expressed genes associated with each cell type.

Figures 5 and 6 compare the true and estimated cell proportions using different methods under the two evaluated datasets. For example, the scatter plot on the first row of Figure 5 represents the deconvolution results when the expression profile of Jurkat cells was unknown to all the methods. According to the characteristics of the semi-reference-based approach, the proportions of Jurkat cells in mixed samples can be recovered through estimating unknown cell proportions. Notably, the constrained models, NNLS and EPIC, tend to assign a very small proportion to the unknown cell component. This phenomenon was observed in the supplementary material of robustness and can be attributed to the loss of the shift-invariant property, which causes the shrinkage of unknown cell proportion estimates.

**Figure 5.**
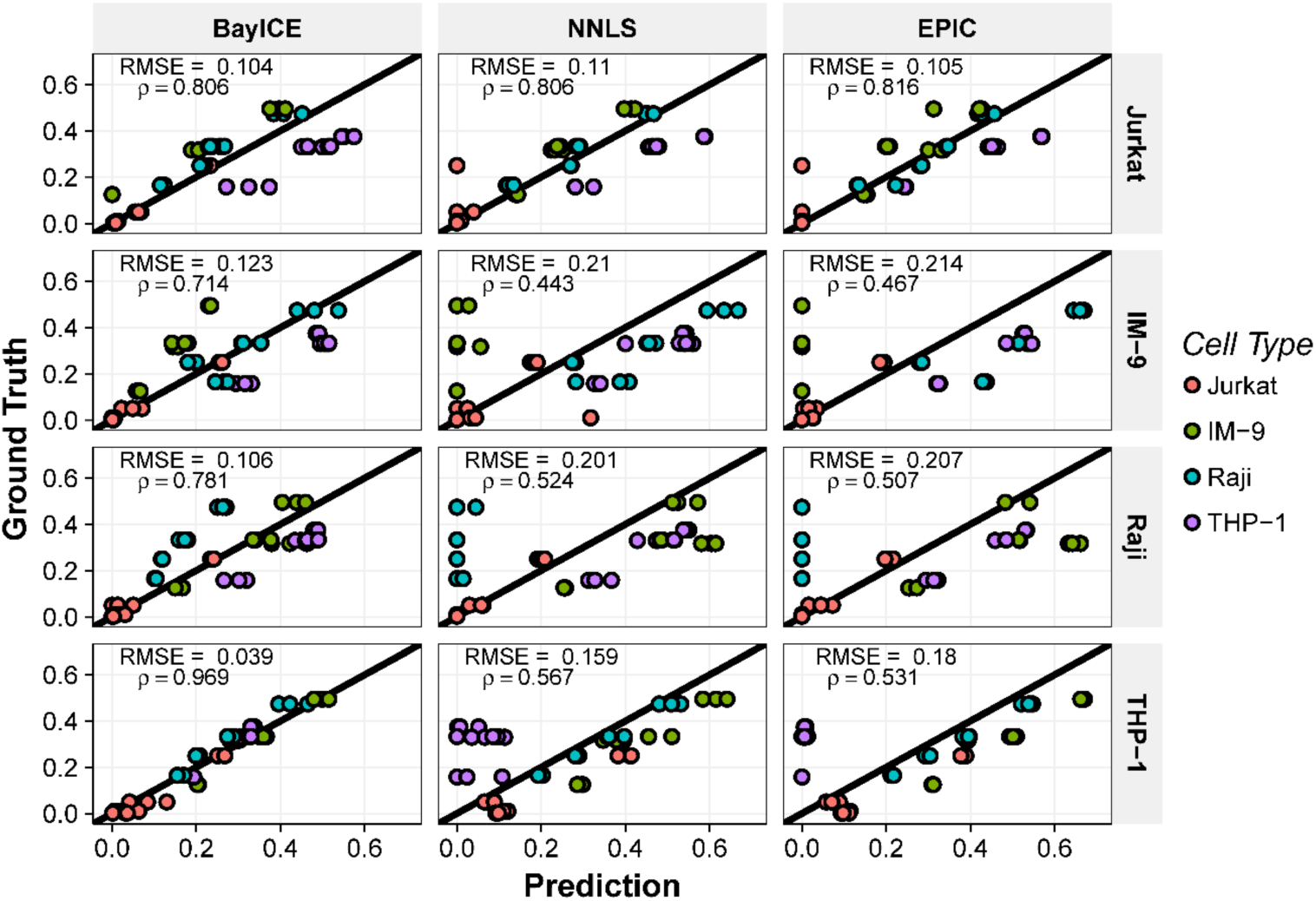
Validation by GSE11058. Scatter plots comparing estimated cell proportions between the true and estimated proportions from the results of GSE11058. Each column represents a particular method of deconvolution. The row name indicates the cell type that was masked from the reference set and referred to as the unknown cell type. Each of the 12 mixed sample results in four estimates of weights and thus 4 points in each plot. The root-mean-square error and correlation between the ground truth and estimation are also provided in the upper-left corner of each plot.

**Figure 6.**
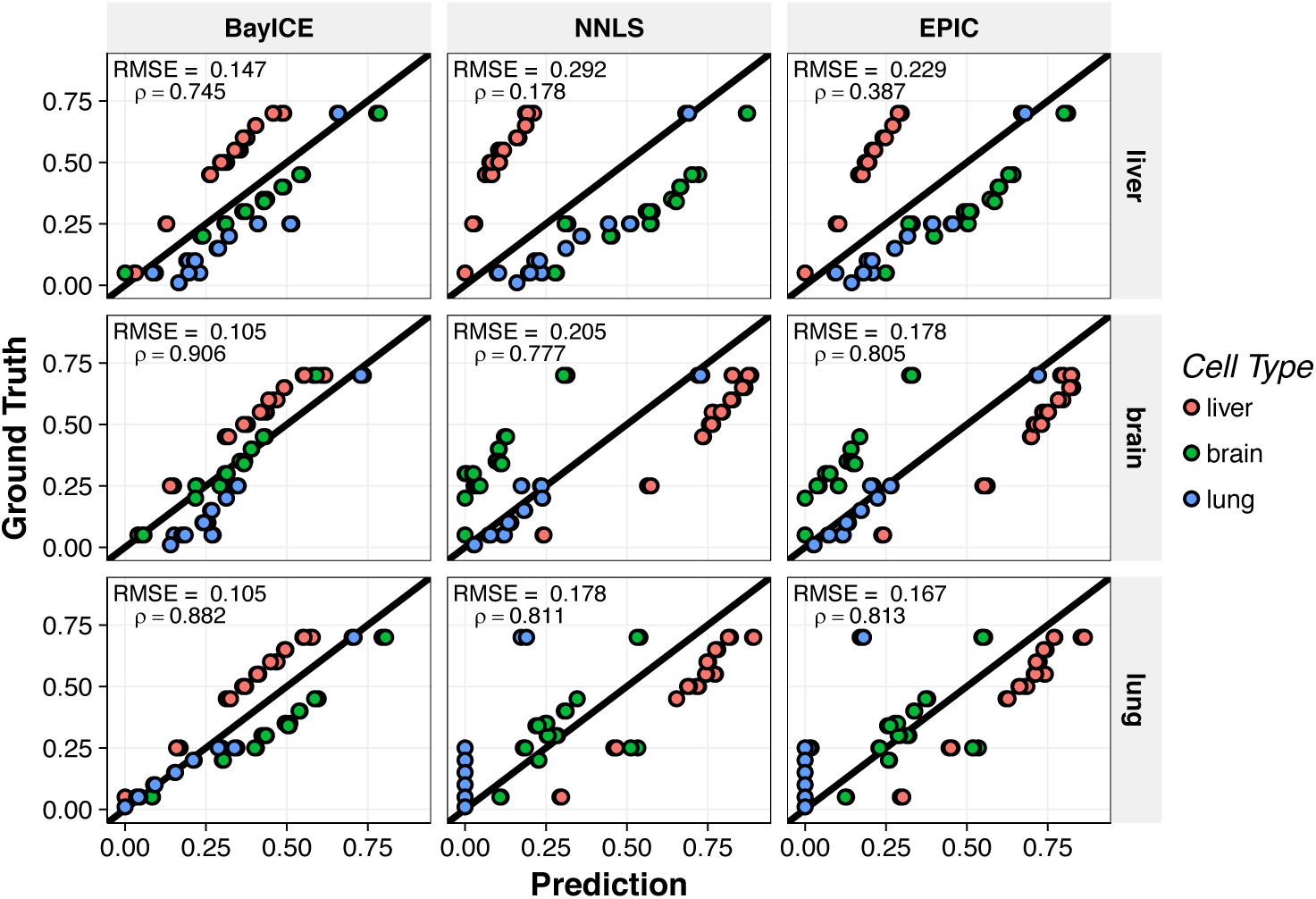
Validation by GSE19830. Scatter plots comparing the true and estimated proportions from the results of GSE19830. Each column represents a particular method of deconvolution. The row name indicates the cell type that was masked from the reference set and referred to as the unknown cell type. Each of the 9 mixed sample results in three estimates of weights and thus three points in each plot. The root-mean-square error and correlation between the ground truth and estimation are also provided in the upper-left corner of each plot.

Furthermore, we used the RMSE and Pearson correlation to assess the performance. RMSE is a measure of accuracy, and the average RMSE of BayICE is 0.093, significantly lower than those of NNLS (0.170) and EPIC (0.176). Similarly, the measurement of correlation value is used to monitor the relative order of cellular proportion estimates, and BayICE also outperformed NNLS and EPIC in terms of the relative size (BayICE = 0.82, NNLS = 0.59, and EPIC = 0.58). Apparently, BayICE possesses the advantage of incomplete data deconvolution. Furthermore, the difficulty of deconvolution increases when the abundance of the unknown cell increases. For instance, in Figure 6, the content of liver cells exceeds 50% of the mixed samples on average, and the deconvolution approaches perform relatively worse when liver cells are excluded from the reference set. The aforementioned observation is consistent with our simulation results.

## 6 Application to Non-Small Cell Lung Cancer

To demonstrate an application of BayICE to a biological problem, we consider RNA-seq data of lung tissues from patients with non-small cell lung cancer (NSCLC). Because tumor-infiltrating lymphocytes play a critical role in cancer treatment, exploring the changes in components of immune cells across tumors is of interest. Thus, we consider 199 patients with NSCLC obtained from GSE81089, and apply BayICE to estimate the cellular components of the tumor-infiltrating lymphocytes in each tumor sample (Mezheyeuski, et al., 2018). To construct the reference set, we collect RNA-seq samples of B cells, T cells, granulocytes, and monocytes of blood tissues from GSE51984 (Pabst, et al., 2016). In addition to immune cell types, we include normal lung tissues in the reference set, and the expression profiles of 19 normal lung tissues from GSE81089 are used to infer the contents of normal lung cells in tumors. Because the immune cells are also present in normal lung tissues, we first apply BayICE to normal lung samples to extract the purified gene expression of lung cells and estimate the immune cell components in normal samples. Additionally, we randomly divided 19 normal samples into two sets: ten samples are used for extracting purified expression of lung cells in Section 6.1, and nine samples are analyzed along with tumors for validation in Section 6.2. As a result, we used the complete reference set consisting of immune cell profiles and purified lung cell profiles to recover the cellular proportions of each NSCLC sample.

### 6.1 Deconvolution of normal lung tissues

Figure 7 illustrates the estimated cellular proportions of ten normal lung samples based on the B cells, T cells, granulocytes, and monocytes. The results of normal lung tissue deconvolution reveal that monocytes are more prevalent than the other immune cells in lung tissues. Monocytes typically circulate through the blood for 1–3 days before migrating into tissues, where they become macrophages or dendritic cells. In lungs, monocytes migrate from the bloodstream into the pulmonary alveolus and are specifically called alveolar macrophages, which play a critical role in homeostasis, host defense, response to foreign substances, and tissue remodeling (Kopf, et al., 2015).

**Figure 7.**
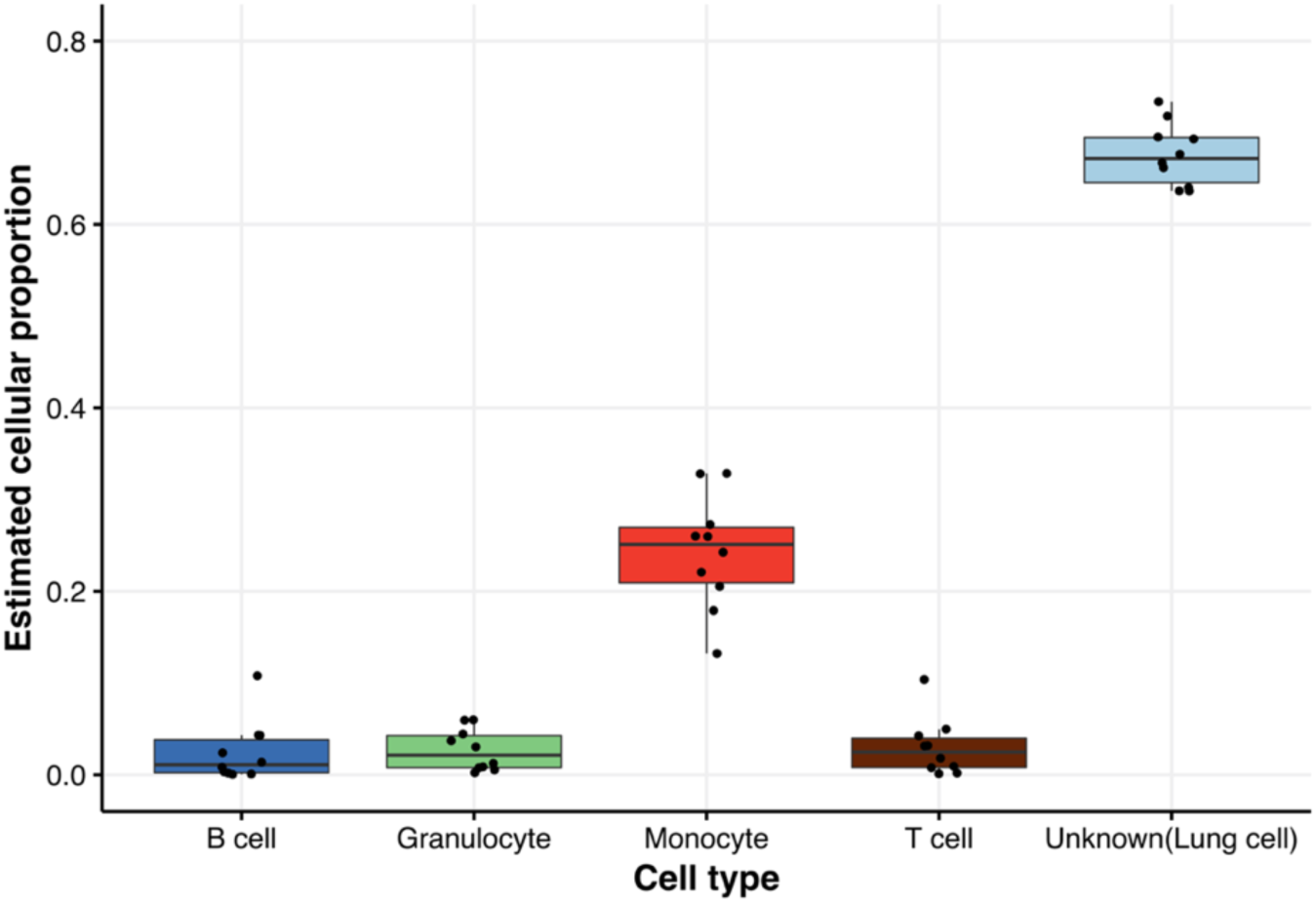
Cell proportion estimation of normal lung tissues. Plot showing estimated proportions of different cell types from 10 normal lung tissues. The rightmost box represents the estimated unknown component from BayICE, and we refer to this component as normal lung cells.

In addition to the immune cell types collected in the reference set, BayICE could extract the component of one uncharacterized cell type present in tissues. In this case, the unknown cell type was presumably dominated by normal lung cells, and we calculated the mean expression profile of the unknown cell type using the posterior mean. The next step was to integrate the immune cell profiles with the estimated profile of normal lung cells to construct a new reference set for tumor sample deconvolution.

### 6.2 Deconvolution of tumors

In this step, we investigated the 199 NSCLC samples and nine healthy samples according to the new reference set. We applied BayICE to these 208 samples and estimated the fractions of normal lung cells, B cells, T cells, granulocytes, and monocytes, as well as malignant cells. Following the mechanism of normal tissue deconvolution, the malignant cell fractions were defined as the unknown cell proportions in tumor deconvolution. To more effectively understand the change in cellular components during tumor progression, we considered the classification of a malignant tumor (TNM) staging system, which is a standard for classifying the extent of cancer spread. Figure 8 is a boxplot of estimated cell proportions under different TNM stages.

**Figure 8.**
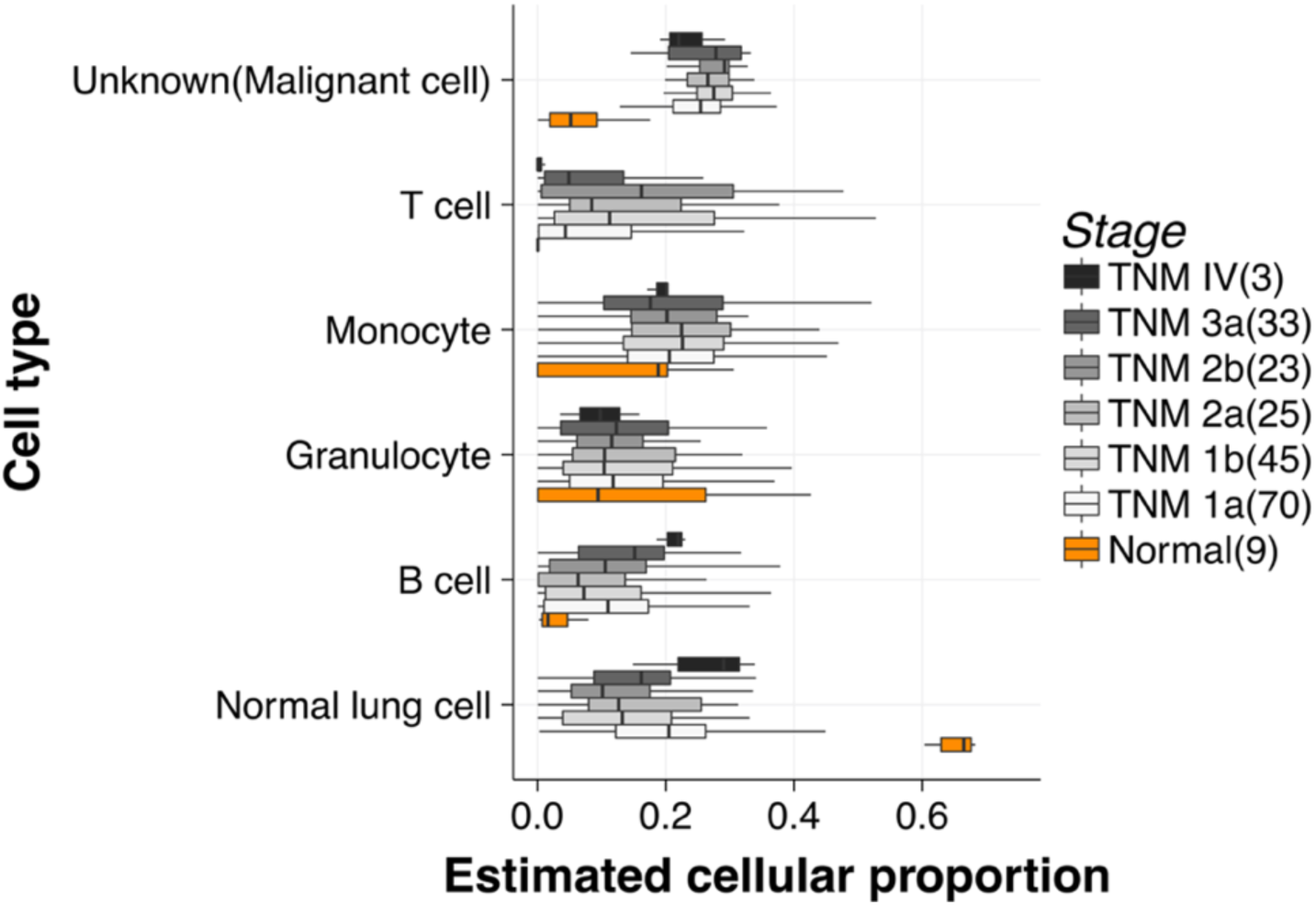
Box plot of cell proportion estimation. Orange boxes refer to the results of the nine normal lung samples, and other boxes are results from 199 patients with non-small cell lung cancer. The tumor samples can be grouped into six stages and the sample number included for each stage is indicated in the parentheses. For samples of the same tumor stage, the estimated proportions of any specific cell type are summarized as one boxplot and the boxplots for the same cell type are plotted side-by-side.

Two crucial observations were made from the deconvolution results. First, the estimated cell proportions were more consistent between healthy samples than between patients with NSCLC. For example, the interquartile ranges of estimated lung cell proportions in normal tissues and tumors were 0.047 and 0.137, respectively. A three-fold difference of dispersion between healthy samples and patients revealed that the homeostatic balance between cells in the lung is disturbed when tissues are cancerous. The high fluctuation of cellular components between patients with NSCLC can directly explain the inter-tumor heterogeneity, which leads to the different attributes of different tumors despite the same diagnoses. Second, our study showed that the number of immune cells in the tissue microenvironment increases from normal stage to cancerous stage, which is in agreement with past studies (Banat, et al., 2015; Li, et al., 2016; Ruffini, et al., 2009; Seo, et al., 2018). In particular, (Banat, et al., 2015) comprehensively assessed the number of immune cells in lung cancer by directly counting cells with cell-specific biomarkers and observed an increased number of immune cells in lung cancer tissues compared with healthy donor lungs. In 2018, Seo et al. (2018) applied two approaches, ESTIMATE and TIMER, to infer the cellular components in NSCLC (Li, et al., 2016; Seo, et al., 2018; Yoshihara, et al., 2013). Their results showed a high abundance of dendritic cells (derived from monocytes), which coincides with our observation of estimated monocyte proportions.

## 7 Discussion and Conclusion

In this study, we developed a novel deconvolution model, BayICE, to infer the cellular components of bulk tissues. BayICE is a semi-reference-based approach that aims to explore cell populations characterized by an external reference set of purified samples and simultaneously investigate the content of uncharacterized cells present in bulk tissues. However, in contrast to constrained models, BayICE takes advantage of a hierarchical Bayesian framework to not only estimate the unknown cell proportion but also recover its gene expression profile. Furthermore, BayICE maintains the shift-invariant property, which guarantees the accuracy of cell proportion estimation.

Other than the above-mentioned merits, there are two major contributions of this study. First, BayICE integrates gene expression deconvolution and gene selection in the same model. Most of the current deconvolution approaches require pre-analysis of an additional dataset to identify signature genes for deconvolution, and the external gene selection might not be consistent to the target dataset. Thus, BayICE incorporates SSVS, a Bayesian variable selection approach, to implement internal gene selection. Second, BayICE adopts shared parameters between the pure cells and tumor samples for cell-specific effect. It has been studied that the cell-to-environment interaction causes some of the genes to be expressed inconsistently after cell sorting. We have shown that the joint modeling of both pure cells and tumors in BayICE is more resistant to the problem of inconsistent genes.

To evaluate the model’s performance, we first conducted an analysis under several simulation scenarios to investigate cell proportion estimation, gene identification, and the robustness of the model. For proportion estimation, we compared two other semi-reference-based approaches, EPIC and NNLS, with BayICE under different unknown cell contents. The results revealed that BayICE significantly outperformed the other methods. We further provided a partial comparison of gene identification with the well-known gene selection approaches, DESeq and edgeR. We found that BayICE can decompose bulk data extremely well and identify cell type-related genes. Moreover, a simulation study using three different simulators revealed that BayICE is more robust with respect to data types. In addition to simulation data, we further applied two real datasets with underlying true cell proportions for validation. The validation of incomplete data deconvolution was consistent with our simulation results, in which BayICE exhibited high accuracy in cell proportion estimation.

Our real data application presented an example using the data of 199 patients with NSCLC. We first applied BayICE to healthy lung samples to investigate the cellular components under a normal condition and extract the relatively purified expression profile of lung cells. The deconvolution of normal tissues succeeded in capturing the primary component of immune cells. Next, we formed a new reference set consisting of the immune cell profiles and estimated normal lung cell profiles to analyze the patients with NSCLC. From the analysis, we observed inter-tumor heterogeneity in NSCLC according to the high variation in cell proportions across tumors. In addition, when comparing immune cell proportions between normal and NSCLC samples, the increased immune cell content in tumors revealed that the immune system was highly activated in the cancerous microenvironment. The inference of NSCLC deconvolution coincides with not only the analytic deconvolution results from other studies but also the observations from an immunohistochemical experiment.

BayICE has thoroughly addressed technical problems of deconvolution to investigate cellular components, but one issue relating to cell activity remains. In real applications, the activities of cell-to-cell communication and cell-to-environment interaction cause some of the genes to be expressed inconsistently between reference and bulk tissue samples. This phenomenon increases the difficulty of selecting marker genes, and an inappropriate gene set limits the ability to explore tissue environments. Although the joint modeling technique of BayICE can adjust the biased gene expression induced by cell sorting, integrating the biological information of cell activity with deconvolution is believed to be more efficient in estimating cell proportions of bulk tissues. Single-cell RNA-seq has emerged as a powerful new set of technologies for characterizing cell interaction, and the primary goal of our future studies is incorporating single-cell RNA data to modify the prior structure with cell interaction. This extension will generate new insights into the deconvolution framework.

Moreover, we will also investigate the unknown cell proportions estimated by BayICE. The proportion of the unknown cell class can be further decomposed if the unknown class is comprised of multiple cancerous cell types. Instead of estimating immune cell proportions, the study of tumor clonal evolution focuses on exploring the contents of different tumor cell types to understand tumor progression. Hence, after BayICE dissects the part belonged to tumor, we can further decompose tumor cells to evaluate the size of tumor clones.

All of the work in this study was implemented on R, and the R package, BayICE, is publicly available for deconvolution analysis (https://github.com/AshTai/BayICE).

## Acknowledgments

This work was supported by the Ministry of Science and Technology [MOST xxxx].

## Notes

https://github.com/AshTai/BayICE

