## Supplementary material for "BayICE: A hierarchical Bayesian deconvolution model with stochastic search variable selection"

**1.	Shift-invariant property**

In this section of the supplementary material, we comprehensively discussed the shift-invariant property for the reference-free, reference-based, and semi-reference-based approaches. Furthermore, we also investigated the shift-invariant property to the proposed Bayesian deconvolution model.

The constrained least square with a complete reference set has an identical estimate against data with different location parameters. To show that, we assume that $y_{g}+c$ and $B_{gk}+c$ are the shifted data with a constant number c, and the proof is given by:

${argmin}_{\left| \mathbf{W} \right|=1}\sum_{g} \left( \left( y_{g}+c \right)-\sum_{k} \left( B_{gk}+c \right)w_{k} \right)^{2}$

$={argmin}_{\left| \mathbf{W} \right|=1}\sum_{g} \left( y_{g}-\sum_{k} B_{gk}w_{k}+c\times\left( 1-\sum_{k} w_{k} \right) \right)^{2}$

$={argmin}_{\left| \mathbf{W} \right|=1}\sum_{g} \left( y_{g}-\sum_{k} B_{gk}w_{k} \right)^{2},$

where the second equality follows by the constraint of $w_{k}$,$\sum_{k} w_{k}=1$. As a result, if the mixed expression ($y_{g}$) and cell type-specific expression ($B_{gk}$) are from the same experiment and have identical mean shift, then the proof above reveals that the reference-based deconvolution approach is shift-invariant. The shift-invariant property also holds in the reference-free approach, because it directly decomposes bulk expression into several components with the identical location parameter.

Nevertheless, the existing semi-reference-based deconvolution methods usually violate shift-invariant property. The current semi-reference-based approaches adopt inequality-constrained optimization by the following formation:

(1.1)

$\hat{\boldsymbol{W}}={argmin}_{\left| \mathbf{W} \right|<1}\sum_{g} \left( y_{g}-\sum_{k} B_{gk}w_{k} \right)^{2}$.

Similarly, we consider the shifted data $y_{g}+c$ and $B_{gk}+c$ to equation (1.1), and we have

(1.2)

${argmin}_{\left| \mathbf{W} \right|<1}\sum_{g} \left( \left( y_{g}+c \right)-\sum_{k} \left( B_{gk}+c \right)w_{k} \right)^{2}$

$={argmin}_{\left| \mathbf{W} \right|<1}\sum_{g} \left( y_{g}-\sum_{k} B_{gk}w_{k}+c\times\left( 1-\sum_{k} w_{k} \right) \right)^{2}$.

Different from the argument in the reference-based approach, the second term $(1-\sum_{k} w_{k})$ in (1.2) is non-zero due to the constraint $\left| \boldsymbol{W} \right|<1$. The presence of the non-zero term $(1-\sum_{k} w_{k})$ implies that the solution to the optimization in (1.2) depends on the amount of c. Additionally, if the value of c increases, then the proportion of unknown cell type $1-\sum_{k} w_{k}$ will shrink toward zero due to minimization. The phenomena of shrinking $1-\sum_{k} w_{k}$ can be seen in our simulation study and analysis of validation data. This fact reveals that inequality-constrained model may be less reliable for exploring unknown cell proportion in deep RNA-seq studies.

In the Bayesian framework, we assume a prior distribution on $B_{gk}$ and denote it as $f(B_{gk};\mu_{gk})$. $\mu_{g}$ is the hyper-parameter of the mean. Under Gaussian assumption, the objective function of Bayesian modeling is

$\sum_{g} \left( y_{g}-\sum_{k} B_{gk}w_{k}+B_{g0}w_{0} \right)^{2}+\sum_{k} \left( B_{gk}-\mu_{g} \right)^{2}+\left( B_{g0}-\mu_{g} \right)^{2}$,

where $\{B_{g0}\}$ is the unknown profile and $w_{0}$ is the unknown proportion. We can then minimize the objective function to obtain the estimates of cell proportions as below:

$$\hat{\boldsymbol{W}}={argmin}_{\left| \mathbf{W} \right|=1}\sum_{g} \left( y_{g}-\sum_{k} B_{gk}w_{k}+B_{g0}w_{0} \right)^{2}+\sum_{k} \left( B_{gk}-\mu_{g} \right)^{2}+\left( B_{g0}-\mu_{g} \right)^{2}.$$

If we plug the shifted data $y_{g}+c$ and $B_{gk}+c$ into the equation above, then we obtain

$${argmin}_{\left| \mathbf{W} \right|=1}\sum_{g} \left( \left( y_{g}+c \right)-\sum_{k} \left( B_{gk}+c \right)w_{k}+\left( B_{g0}+c \right)w_{0} \right)^{2}+\sum_{k} \left( \left( B_{gk}+c \right)-\mu_{g} \right)^{2}+\left( \left( B_{g0}+c \right)-\mu_{g} \right)^{2}$$

$$={argmin}_{\left| \mathbf{W} \right|=1}\sum_{g} \left( y_{g}-\sum_{k} B_{gk}w_{k}+B_{g0}w_{0}+c(1-\sum_{k} w_{k}-w_{0}) \right)^{2}+\sum_{k} \left( B_{gk}-\left( \mu_{g}-c \right) \right)^{2}+\left( B_{g0}-\left( \mu_{g}-c \right) \right)^{2}$$

Because of the constraint $1-\sum_{k} w_{k}-w_{0}=0$, the estimate obtained from the equation above is exactly equal to $\hat{\boldsymbol{W}}$. It reveals that the Bayesian deconvolution method follows the shift-invariant property.

**2.	Normalization**

*TPM approach*

Calculating TPM includes the following three standard steps.

1. Divide the read counts by the length of each gene in kilobases. It is the *reads per kilobase* (RPK).
2. Count up all the RPK values in an individual and divide this number by 1,000,000. That is the “per million” scaling factor.
3. Divide the RPK values by the “per million” scaling factor, and then the result is the TPM.

*Resampling approach*

Let $E_{gi}$ be the read count of gene *g* for sample *i*. The resampling approach is conducted as follows:

1. Calculate total read count of sample *i* and denote it as $T_{i}=\sum_{g} E_{gi}$.
2. Resample the gene expression $E_{gi}^{*}$ by Binomial distribution as

$E_{gi}^{*}\sim Bin(E_{gi},\frac{\min\left( T_{i} \right)}{T_{i}}P )$,

where P is an adjusted number

This procedure guarantees that the resampled total read count of every samples are approximately the same as $\min\left( T_{i} \right)P$.

**3.	The Gibbs sampler in BayICE works**

The Gibbs sampler in BayICE works as follows:

1. Conditional on all other parameters, $\alpha_{g}$ is simulated from:

$$\left( \alpha_{g} | . \right) \sim N\left( a_{g},A_{g} \right),$$

where $a_{g}=\frac{\left( \eta_{g}^{2} \right)^{-1}\sum_{i} \left( y_{gi}^{*}-\sum_{k'} x_{ik'}\beta_{gk'} \right)+M\left( \tau_{g}^{2} \right)^{-1}\sum_{j} \left( z_{gj}^{*}-\sum_{k'} w_{jk'}\beta_{gk'} \right)}{N\left( \eta_{g}^{2} \right)^{-1}+M\left( \tau_{g}^{2} \right)^{-1}}$ and

$A_{g}=\left( \frac{N}{\left( N+M \right)\eta_{g}^{2}}+\frac{M}{\left( N+M \right)\tau_{g}^{2}} \right)^{-1}$.

2. Conditional on all other parameters, $\beta_{gk}$ is simulated from:

$$\left( \beta_{gk} | . \right) \sim N\left( b_{gk},B_{gk} \right),$$

where

$$b_{gk}=\frac{\frac{\sum_{i} \left( y_{gi}^{*}-\alpha_{g}-\sum_{k^{'}\neq k} x_{ik^{'}}\beta_{gk^{'}} \right)}{\sum_{i} x_{ik}}\times r1+\frac{\sum_{j} \left( z_{gj}^{*}-\sum_{k^{'}\neq k} w_{jk^{'}}\beta_{gk^{'}} \right)w_{jk}}{\sum_{j} \left( w_{jk} \right)^{2}}\times r2}{r1+r2+r3},$$

$$r1=\left( \frac{\left( N+M \right)\eta_{g}^{2}}{\sum_{i} \sum_{k^{'}\neq k} x_{ik^{'}}} \right)^{-1}, r2=\left( \frac{\left( N+M \right)\tau_{g}^{2}}{\left( \sum_{j} \sum_{k^{'}\neq k} w_{jk^{'}} \right)^{2}} \right)^{-1},$$

$$r3=\left( \psi_{gk}^{2}t_{gk}\tau_{g}^{2} \right)^{-1},$$

and

$$B_{gk}=\frac{1}{r1+r2+r3}.$$

3. Conditional on all other parameters, $\beta_{g0}$ is simulated from:

$$\left( \beta_{g0} | . \right) \sim N\left( b_{g0},B_{g0} \right),$$

where

$$b_{gT}=\frac{\frac{\sum_{j} \left( z_{gj}^{*}-\sum_{k^{'}\neq0} w_{jk^{'}}\beta_{gk^{'}} \right)w_{j0}}{\sum_{j} \left( w_{j0} \right)^{2}}\times s1}{s1+s2},$$

$$s1=\left( \frac{\left( N+M \right)\tau_{g}^{2}}{\left( \sum_{j} \sum_{k^{'}\neq0} w_{jk^{'}} \right)^{2}} \right)^{-1}, s2=\left( \psi_{gk}^{2}t_{g0}\tau_{g}^{2} \right)^{-1},$$

and

$$B_{g0}=\frac{1}{s1+s2}.$$

4. Conditional on all other parameters, $\eta_{g}^{2}$ is simulated from:

$$\eta_{g}^{2} \sim Inv\Gamma\left( \frac{N}{2}-1,\frac{RSS.ref.g}{2} \right),$$

where

$$RSS.ref.g=\frac{\sum_{i} \left( y_{gi}^{*}-\alpha_{g}-\sum_{k^{'}} x_{ik^{'}}\beta_{gk^{'}} \right)^{2}}{N+M}.$$

5. Conditional on all other parameters, $\tau_{g}^{2}$ is simulated from:

$$\tau_{g}^{2} \sim Inv\Gamma\left( \frac{M+(K+1)}{2}-1,\frac{RSS.mix.g+RSS.beta.g}{2} \right),$$

where

$$RSS.mix.g=\frac{\sum_{i} \left( y_{gi}^{*}-\alpha_{g}-\sum_{k^{'}} w_{jk^{'}}\beta_{gk^{'}} \right)^{2}}{N+M},$$

and

$$RSS.beta.g=\frac{\sum_{k^{'}} \left( \frac{\left( \beta_{gk^{'}} \right)^{2}}{\psi_{gk}^{2}t_{gk}} \right)}{N+M}.$$

6. Conditional on all other parameters, $\psi_{gk}^{2}$ is simulated from:

$$\psi_{gk}^{2} \sim Inv\Gamma\left( a_{0}+\frac{1}{2},b_{0}+\frac{\left( \beta_{gk} \right)^{2}}{t_{gk}\tau_{g}^{2}} \right).$$

7. Conditional on all other parameters, $t_{gk}$ is simulated from:

$$t_{gk} \sim\frac{u_{gk1}}{u_{gk1}+u_{gk2}}I_{1}\left( t_{gk} \right)+\frac{u_{gk2}}{u_{gk1}+u_{gk2}}I_{c}\left( t_{gk} \right),$$

where

$$u_{gk1}=\pi_{k}\exp\left( -\frac{\left( \beta_{gk} \right)^{2}}{2\psi_{gk}^{2}\tau_{g}^{2}} \right),$$

and

$$u_{gk2}=\left( 1-\pi_{k} \right)c^{-\frac{1}{2}}\exp\left( -\frac{\left( \beta_{gk} \right)^{2}}{2c\psi_{gk}^{2}\tau_{g}^{2}} \right).$$

8. It simulates $\pi_{k}$ from its conditional distribution,

$$Beta\left( c_{0}+\sum_{k} I\left( t_{gk}=1 \right) ,d_{0}+\sum_{k} I\left( t_{gk}=c \right) \right).$$

9. It simulates the cellular components $w_{jK}$ through the Metropolis–Hastings approach.

**4.	Simulators**

Herein, we introduce the two remaining simulators discussed in the manuscript for the simulation study, namely the normal simulator and negative binomial (NB) simulator. The notations in this document are consistent with the manuscript. All of the parameters for the normal simulator and NB simulator are generated using the same generation rule as the multinomial generator. The normalized effect size is fixed at 0.5. The complete sampling procedures of the normal simulator and NB simulator are presented in Algorithms 1 and 2, respectively. The first five steps of the three simulators are identical, and the last two steps are simulator-specific procedures.

*Sampling procedure of multinomial generator*

Step 1: Randomly pick 5000 genes of 17,775 genes from GSE81089 and calculate their mean expression $\left\{ a_{g}^{sim} \right\}$.

Step 2: Generate $\left\{ c_{g}^{sim} \right\}$ from Ber(0.5) and then calculate the cell-specific effect $\left\{ b_{g}^{sim} \right\}$.

Step 3: Generate $\{e_{ref,i}^{sim,}\}$ and $\{e_{mix,j}^{sim}\}$ from $Uniform(0.9,1.1)$.

Step 4: Set $w_{j0}^{sim}$ as one of $\{0.1, 0.2,\ldots, 0.9\}$ and for each of the $w_{j0}^{sim}$ values, generate $\left( w_{j1}^{sim},w_{j2}^{sim},w_{j3}^{sim},w_{j4}^{sim} \right)$ from Dirichlet(1,…,1).

Step 5: Normalize $(w_{j1}^{sim},\ldots,w_{j4}^{sim},w_{j0}^{sim})$ as $(w_{j1}^{*sim},\ldots,w_{j4}^{*sim},w_{j0}^{*sim})$ by

$$w_{jk}^{*sim}=\frac{w_{jk}^{sim}}{1+w_{j0}^{sim}} \mathrm{and} w_{j0}^{*sim}=\frac{w_{j0}^{sim}}{1+w_{j0}^{sim}}.$$

Step 6: Generate the expression profile of sample *i* in the reference set

$$\left( y_{1,i}^{sim},\ldots,y_{5000,i}^{sim} \right)\sim Multinomial(N=5\times{10}^{6},p=(p_{1,i}^{sim},\ldots,p_{5000,i}^{sim}))$$

where $\left( p_{1,i}^{sim},\ldots,p_{5000,i}^{sim} \right)\propto\left( e_{ref,i}^{sim,}b_{1}^{sim}+\sum_{k=1}^{4} x_{ik}b_{1}^{sim},\ldots,e_{ref,i}^{sim,}b_{5000}^{sim}+\sum_{k=1}^{4} x_{ik}b_{5000}^{sim} \right)$.

Step 7: Generate the expression profile of sample *j* in the bulk sample set

$$\left( z_{1,j}^{sim},\ldots,z_{5000,j}^{sim} \right)\sim Multinomial(N=5\times{10}^{6},p=(p_{1,j}^{sim},\ldots,p_{5000,j}^{sim}))$$

where

$$\left( p_{1,j}^{sim},\ldots,p_{5000,j}^{sim} \right)\propto\left( e_{mix,j}^{sim}b_{1}^{sim}+\sum_{k=1}^{4} w_{jk}^{*sim}b_{1}^{sim}+w_{j0}^{*sim}b_{1}^{sim},\ldots,e_{mix,j}^{sim}b_{5000}^{sim}+\sum_{k=1}^{4} w_{jk}^{*sim}b_{5000}^{sim}++w_{j0}^{*sim}b_{5000}^{sim} \right).$$

*Simulation results from the normal simulator and NB simulator*

We applied BayICE, non-negative least-squares (NNLS), and EPIC to two artificial datasets generated by the NB simulator and the normal simulator. The results of recovering unknown cell profiles and estimating cell proportions, which were identical to the summary of the results from the multinomial simulator presented in the manuscript, are illustrated in Figures 1 and 2 of the manuscript. In these figures, two measurements, the root-mean-square error (RMSE) and Pearson correlation, are used to quantify the behavior of BayICE, EPIC, and NNLS in cell proportion estimation. BayICE exhibits the ability to recover the gene expression profile of an unknown cell type, and the comparison results reveal that BayICE outperforms the other approaches under each simulator.

Algorithm 1: NB simulator

**Step 1:** Randomly pick 5000 genes and calculate their mean expression $\left\{ a_{g}^{sim} \right\}$.

**Step 2:** Generate $\left\{ c_{g}^{sim} \right\}$ from Ber(0.5) and then calculate the cell-specific effect $\left\{ b_{g}^{sim} \right\}$.

**Step 3:** Generate $\{e_{ref,i}^{sim,}\}$ and $\{e_{mix,j}^{sim}\}$ from $Uniform(0.9,1.1)$.

**Step 4:** Set $w_{j0}^{sim}$ as one of $\{0.1, 0.2,\ldots, 0.9\}$, and for each of the $w_{j0}^{sim}$ values, generate $\left( w_{j1}^{sim},\ldots,w_{j4}^{sim} \right)$ from Dirichlet(1,…,1).

**Step 5:** Normalize $(w_{j1}^{sim},\ldots,w_{j4}^{sim},w_{j0}^{sim})$ as $(w_{j1}^{*sim},\ldots,w_{j4}^{*sim},w_{j0}^{*sim})$ by

$$w_{jk}^{*sim}=\frac{w_{jk}^{sim}}{1+w_{j0}^{sim}} \mathrm{and} w_{j0}^{*sim}=\frac{w_{j0}^{sim}}{1+w_{j0}^{sim}}.$$

**Step 6:** Generate the expression value of gene *g* of sample *i* in the reference set

$$y_{g,i}^{sim}\sim NB(mean=\mu_{g,i}^{sim},dispersion=1)$$

where $\mu_{g,i}^{sim}=e_{ref,i}^{sim,}b_{g}^{sim}+\sum_{k=1}^{4} x_{ik}b_{g}^{sim}$.

**Step 7:** Generate the expression value of gene *g* of sample *j* in the bulk sample set

$$z_{g,j}^{sim}\sim NB(mean=\mu_{g,j}^{sim},dispersion=1)$$

where $\mu_{g,j}^{sim}=e_{mix,j}^{sim}b_{g}^{sim}+\sum_{k=1}^{4} w_{jk}^{*sim}b_{g}^{sim}++w_{j0}^{*sim}b_{g}^{sim}$.

Algorithm 2: Normal simulator

**Step 1:** Randomly pick 5000 genes and calculate their mean expression $\left\{ a_{g}^{sim} \right\}$.

**Step 2:** Generate $\left\{ c_{g}^{sim} \right\}$ from Ber(0.5) and then calculate the cell-specific effect $\left\{ b_{g}^{sim} \right\}$.

**Step 3:** Generate $\{e_{ref,i}^{sim,}\}$ and $\{e_{mix,j}^{sim}\}$ from $Uniform(0.9,1.1)$.

**Step 4:** Set $w_{j0}^{sim}$ as one of $\{0.1, 0.2,\ldots, 0.9\}$, and for each of the $w_{j0}^{sim}$ values, generate $\left( w_{j1}^{sim},\ldots,w_{j4}^{sim} \right)$ from Dirichlet(1,…,1).

**Step 5:** Normalize $(w_{j1}^{sim},\ldots,w_{j4}^{sim},w_{j0}^{sim})$ as $(w_{j1}^{*sim},\ldots,w_{j4}^{*sim},w_{j0}^{*sim})$ by

$$w_{jk}^{*sim}=\frac{w_{jk}^{sim}}{1+w_{j0}^{sim}} \mathrm{and} w_{j0}^{*sim}=\frac{w_{j0}^{sim}}{1+w_{j0}^{sim}}.$$

**Step 6:** Generate the expression value of gene *g* of sample *i* in the reference set

$$y_{g,i}^{sim}\sim Normal(mean=\mu_{g,i}^{sim},sd=\mu_{g,i}^{sim}/10)$$

where $\mu_{g,i}^{sim}=e_{ref,i}^{sim,}b_{g}^{sim}+\sum_{k=1}^{4} x_{ik}b_{g}^{sim}$.

**Step 7:** Generate the expression value of gene *g* of sample *j* in the bulk sample set

$$z_{g,j}^{sim}\sim Normal(mean=\mu_{g,j}^{sim},sd=\mu_{g,j}^{sim}/10)$$

where $\mu_{g,j}^{sim}=e_{mix,j}^{sim}b_{g}^{sim}+\sum_{k=1}^{4} w_{jk}^{*sim}b_{g}^{sim}++w_{j0}^{*sim}b_{g}^{sim}$.

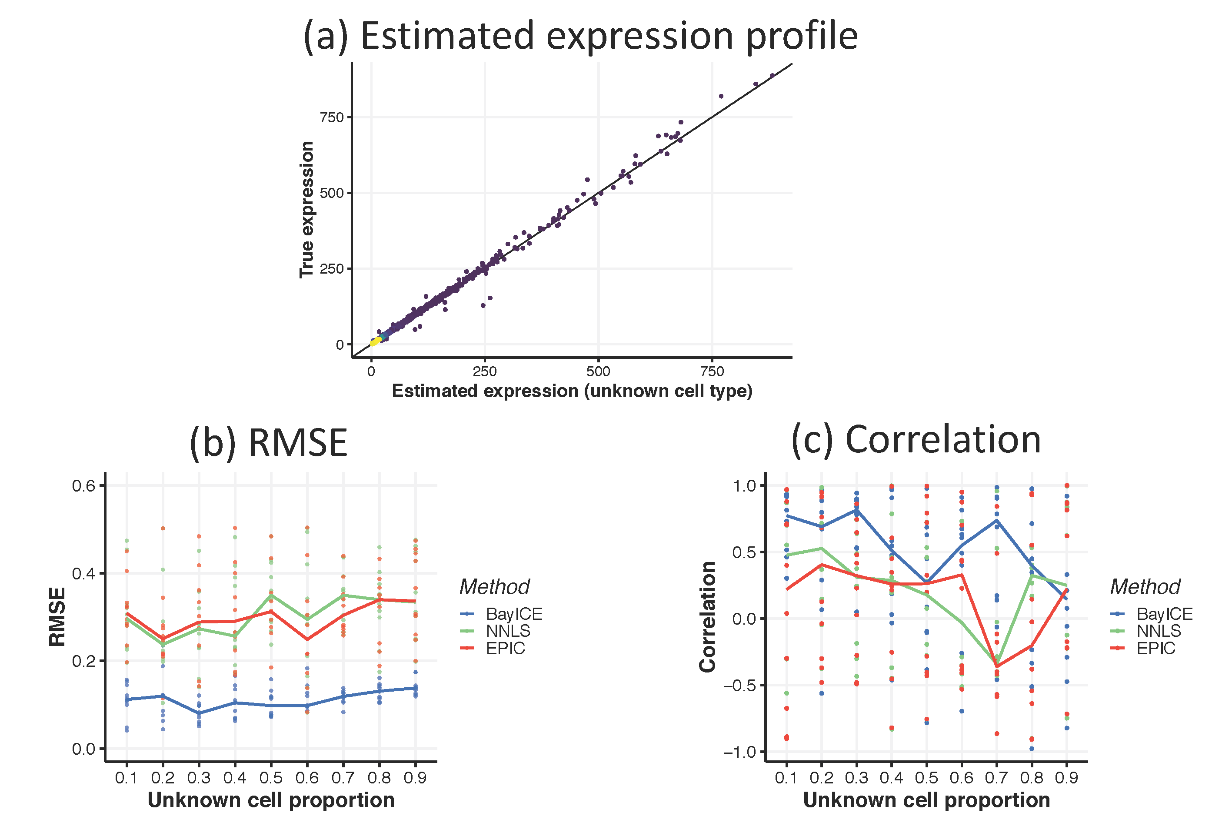

**Figure 1. Deconvolution results from the NB simulator.**

Expression data generated from the NB simulator. (a) Scatter plot of gene expression of the unknown cell type between the truth and estimation. (b) Root-mean-square error between the true and estimated cellular proportions under different levels of unknown cells. Each condition generates 10 random sets. The medians of the 10 random sets are connected as the line in the figure. (c) The correlation between the true and estimated cellular proportions. The median lines are also plotted for comparison.

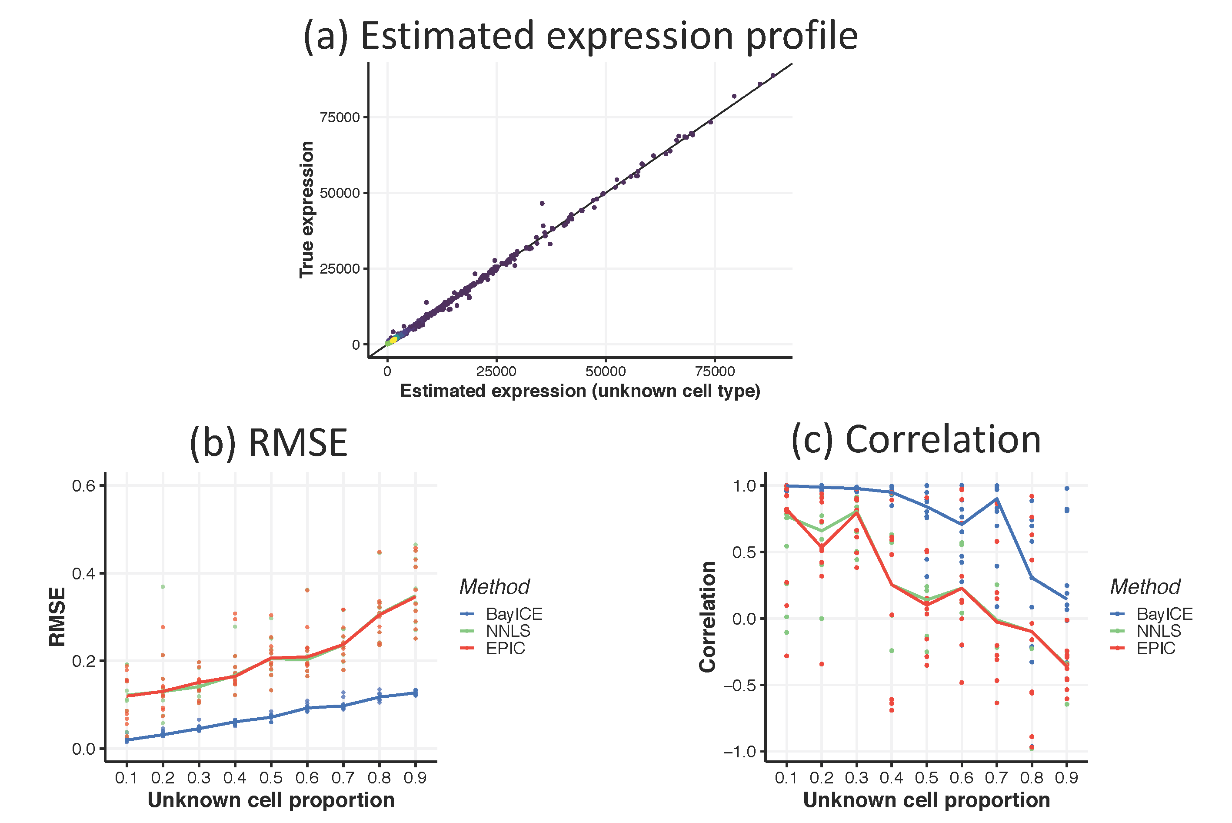

**Figure 2. Deconvolution results from the normal simulator.**

Expression data generated from the normal simulator. (a) Scatter plot of the gene expression of the unknown cell type between the truth and estimation. (b) Root-mean-square error between the true and estimated cellular proportions under different levels of unknown cells. Each condition generates 10 random sets. The medians of the 10 random sets are connected as the line in the figure. (c) The correlation between the true and estimated cellular proportions. The median lines are also plotted for comparison.

**5.	Convergence of MCMC in BayICE**

To demonstrate the convergence property of BayICE, Figure 3 illustrates three trace plots of cell proportions in non-small cell lung cancer study. Normal lung cell, Monocyte, and cancer cell are considered for demonstration. It performs MCMC with 100,000 iterations. It is clear that all estimates of cell proportions converge after 60,000 iterations.

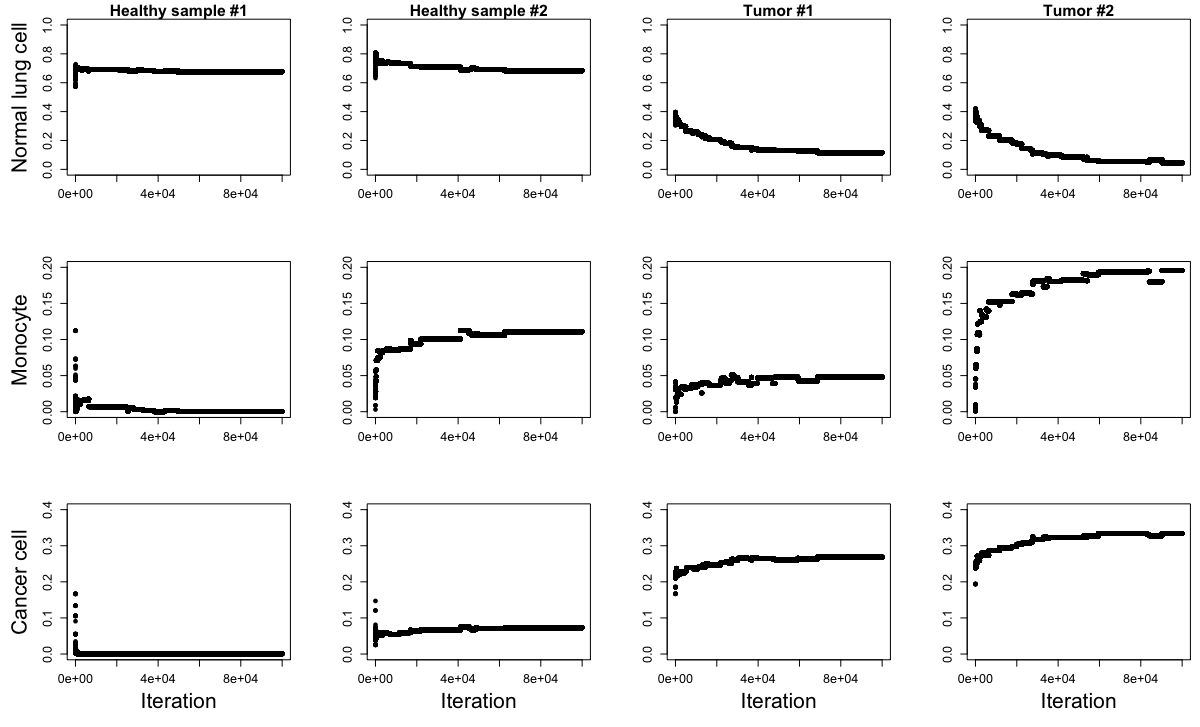

**Figure 3. Trace plot of cell proportions.**

Row indicates cell type and column indicates sample. Each trace plot records 100,000 iterations of MCMC.

**6.	Robustness for deconvolution**

Finally, we examined the robustness of BayICE with respect to different simulators. Following the settings of the multinomial simulator, the normal simulator and negative binomial simulator assume that five cell types are present in tissues and one of the five types is not observed in the reference set. Ninety samples are generated for each simulation in the setting with nine unknown proportions which are 0.1, 0.2, …, 0.8, 0.9. We maintain the same mean structures, parameter settings, and values of cell proportions to generate data based on the real expression profiles from GSE81089. The settings of these simulators and procedures of data generation are shown in the chapter 4 of this file.

We apply BayICE, NNLS, and EPIC to these simulated data to obtain estimations of the cellular components. Since we set the same cell proportions in all simulators, the deconvolution results from analyzing these simulated data can be compared. The results of estimation under three simulators and three approaches are illustrated in Figure 4 of the manuscript. Compared to others, BayICE is relatively unbiased under different simulators. It indicates that BayICE is robust against the distributions of gene expression although the BayICE approach is a Gaussian-based model. By contrast, other approaches exhibit much lower accuracy on the proportion estimation. The main reason to biased proportions estimated by EPIC and NNLS is the lack of shift-invariant property, and it implies the estimates of unknown cell proportions from EPIC and NNLS are biased towards zero. Clearly, BayICE can more effectively adjust for data types compared with the other approaches.

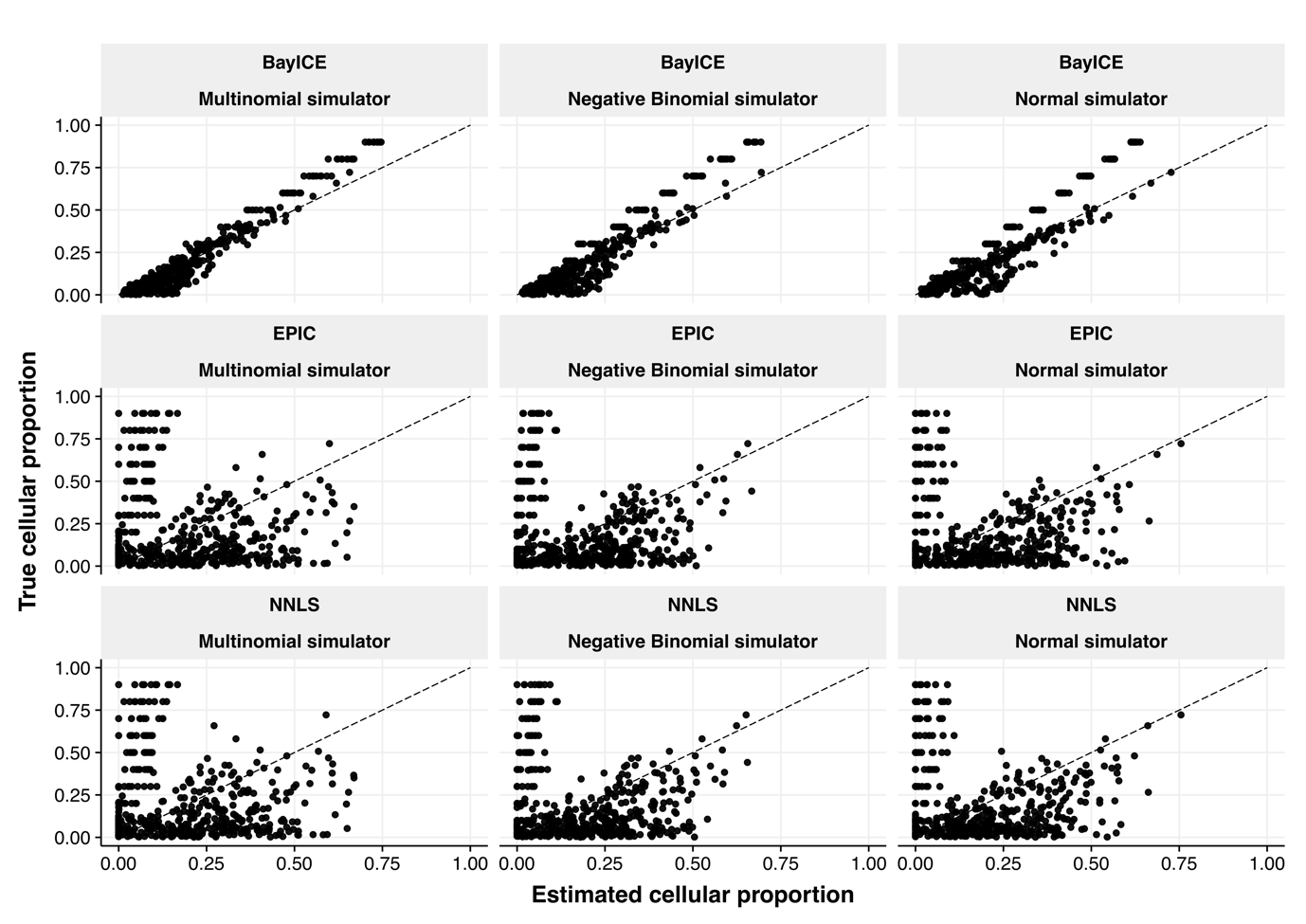

**Figure 4. Scatter plot of the estimation results from data of different simulators.**

Scatter plots between the estimated cell proportions and the true proportions from three simulated datasets. Each row represents a particular method of deconvolution, and each column uses the same data simulated by a particular simulator.
